# Multimodal analyses of immune cells during bone repair identify macrophages as a therapeutic target in musculoskeletal trauma

**DOI:** 10.1101/2024.04.29.591608

**Authors:** Yasmine Hachemi, Simon Perrin, Maria Ethel, Anais Julien, Julia Vettese, Blandine Geisler, Christian Göritz, Céline Colnot

**Affiliations:** Univ Paris Est Creteil, INSERM, IMRB, Creteil, France; Department of Cell and Molecular Biology, Karolinska Institutet, Stockholm, Sweden

**Author notes:** These authors contributed equally (Yasmine Hachemi, Simon Perrin). Corresponding author: Céline Colnot.

## Abstract

Musculoskeletal traumatic injuries (MTI) involve soft tissue lesions adjacent to a bone fracture leading to fibrous nonunion. The impact of MTI on the inflammatory response to fracture and on the immunomodulation of skeletal stem/progenitor cells (SSPCs) remains unknown. Here, we used single cell transcriptomic analyses to describe the immune cell dynamics after bone fracture and identified distinct macrophage subsets with successive pro-inflammatory, pro-repair and anti-inflammatory profiles. Concurrently, SSPCs transition via a pro- and anti-inflammatory fibrogenic phase of differentiation prior to osteochondrogenic differentiation. In a preclinical MTI mouse model, the injury response of immune cells and SSPCs is disrupted leading to a prolonged pro-inflammatory phase and delayed resolution of inflammation. Macrophage depletion improves bone regeneration in MTI demonstrating macrophage involvement in fibrous nonunion. Finally, pharmacological inhibition of macrophages using the CSF1R inhibitor Pexidartinib ameliorates healing. These findings reveal the coordinated immune response of macrophages and skeletal stem/progenitor cells as driver of bone healing and as a primary target for the treatment of trauma-associated fibrosis.

**Summary:** Hachemi et al. report the immune cell atlas of bone repair revealing macrophages as pro-fibrotic regulators and a therapeutic target for musculoskeletal regeneration. Genetic depletion or pharmacological inhibition of macrophages improves bone healing in musculoskeletal trauma.

## Introduction

Musculoskeletal conditions affect approximately 1.71 billion people and are the principal contributor to disability worldwide^1^. Among them, musculoskeletal traumatic injuries (MTI), marked by extensive soft tissue damage associated with a bone fracture, are a leading cause of bone fracture nonunion. Clinical management of MTI is challenging as current treatments are mostly surgical with variable healing outcomes^2^. The consequences of MTI on bone regeneration are poorly understood, limiting our ability to address these debilitating conditions of fracture nonunion.

As observed in many tissues, bone repair following injury is initiated by an inflammatory response and a transient fibrotic response of skeletal stem/progenitor cells (SSPCs), followed by a reparative phase to reestablish tissue integrity and function^3–6^. However, how immune cell dynamics influence the fate of SSPCs after bone injury is still poorly understood. SSPCs reside mostly in the periosteum, as well as bone marrow and skeletal muscles neighboring the fracture site^3,7–11^. Upon fracture, SSPC differentiation occurs in a complex injury environment, where inflammatory cells migrate and play essential roles in the initiation and progression of the repair process. Among inflammatory cell types, macrophages are suspected to be crucial regulators of the early stages of bone healing^12–14^. Inflammatory and resident macrophages remove necrotic cells and debris and secrete chemotactic mediators to recruit SSPCs at the fracture site^12,15–19^. Conversely, prolonged inflammation and delay in the clearance of macrophages may delay healing^20–25^. Specific pro-fibrotic macrophage sub-types have been incriminated in fibrotic diseases and impaired tissue regeneration^19,26–28^. In MTI, the absence of fracture consolidation has been associated with fibrotic accumulation^8^, but the impact of the trauma environment and macrophages on altered SSPC differentiation is still unknown.

Here, we used single-nuclei transcriptomics to describe the dynamic changes of inflammatory cells in response to bone fracture and their relationship with SSPCs. We uncover the heterogeneity of inflammatory cells after bone injury and identified four macrophage subpopulations with distinct signaling roles. We found that SSPCs follow a parallel temporal trajectory with an early pro-inflammatory phase followed by an anti-inflammatory/pro-repair phase. We analyzed a murine model of MTI, where altered SSPC differentiation causes fibrotic accumulation and fracture nonunion^8^. In this MTI model, we showed that the inflammatory response of immune cells and SSPCs is altered, causing fibrous nonunion. Genetic depletion of macrophages improved bone union in the MTI mouse model, revealing macrophages as a promising therapeutic target in MTI. Finally, we showed that the macrophage-specific CSF1R inhibitor can rescue MTI-induced nonunion. More generally, our results highlight the critical role of macrophages in controlling SSPC fate after injury and the promising potential of targeting macrophages to prevent fibrous nonunion.

## Results

### Immune cell heterogeneity in response to bone fracture

To investigate the inflammatory phase of bone repair, we generated a single-nuclei RNAseq dataset of the periosteum and hematoma at day 1 post-fracture. We integrated this dataset with datasets from Perrin *et al.*^4^ containing the uninjured periosteum and the periosteum and hematoma/callus at days 3, 5, and 7 post-fracture (Fig. 1a). We obtained 23 cell clusters corresponding to 11 cell populations i.e. SSPCs, injury-induced fibrogenic cells (IIFCs), osteoblasts, chondrocytes, adipocytes, Schwann cells, fibroblasts, osteoclasts, immune cells, pericytes and endothelial cells (Fig. 1b, Fig. S1a). The immune cell population represented 30% of the dataset, with a peak in the percentage of immune cells at day 1 and 3 post-fracture before a progressive decrease at days 5 and 7 post-fracture (Fig. 1c-d Fig. S1b). We isolated the immune cells and obtained a subset dataset containing 10 cell clusters and corresponding to 6 cell populations: macrophages (expressing *Adgre1* (F4/80)), monocytes (expressing *Cd52*), dendritic cells (expressing *Traf1*), neutrophils (expressing *S100a8*), NK/T cells (expressing *Cd247*) and B cells (expressing *Ighm*) (Fig. 1e, Fig. S1c). The macrophage population was divided in 4 distinct sub-populations: early macrophages were present mostly at day 1 post-fracture and marked by high expression of *Arg1,* intermediate macrophages were present mostly at day 3 post-fracture and marked by high expression of *Cd68,* late macrophages 1 were found from day 5 post-fracture and expressed *Mrc1* (CD206) and late macrophages 2 were found from day 3 and expressed Taco1 (Fig. 1f-g, Fig. S1d-e).

**Figure 1:**
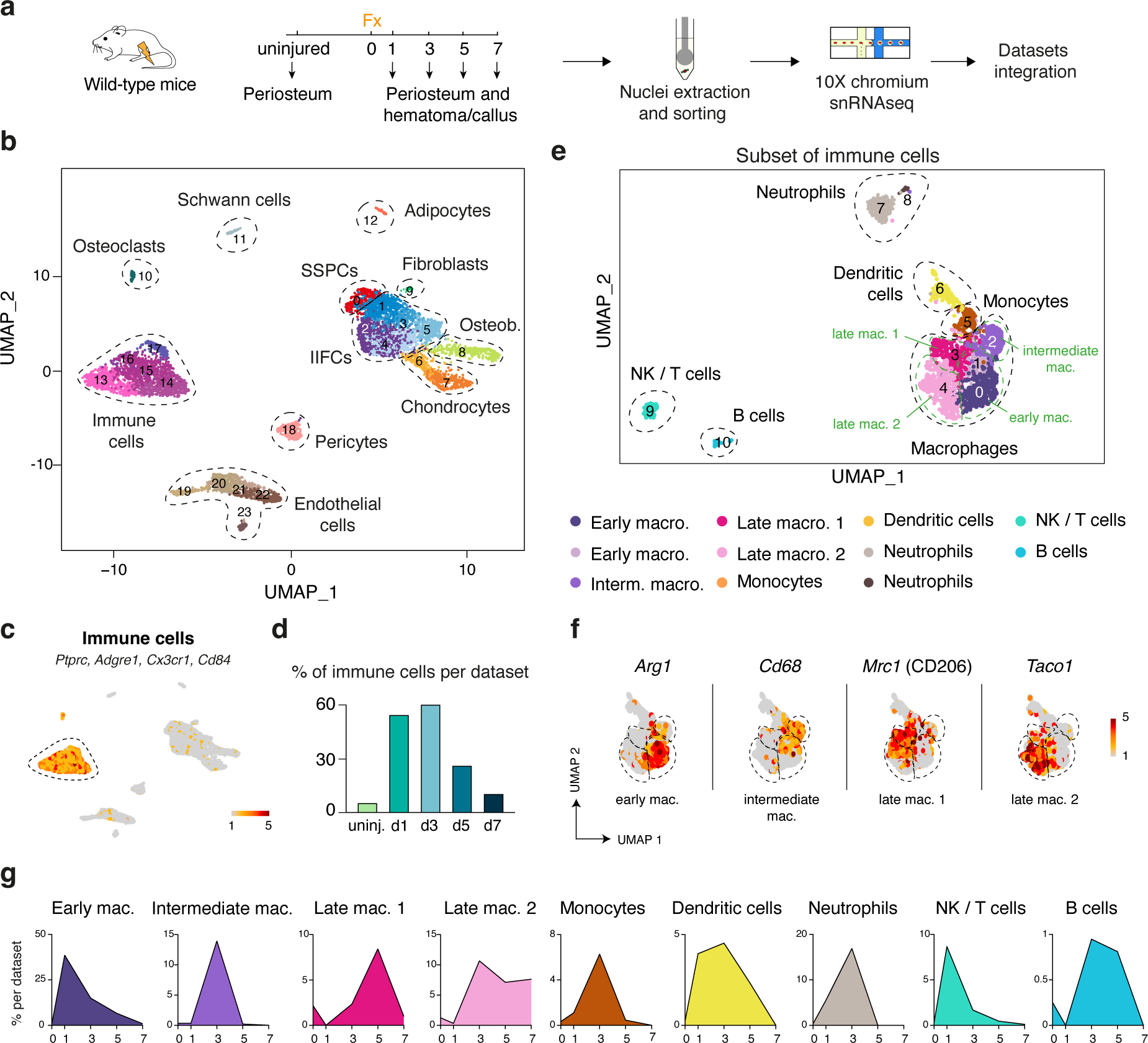
The immune cell atlas of bone regeneration. **a.** Experimental design. Nuclei were extracted from the periosteum of uninjured tibia and from the periosteum and hematoma/callus at days 1, 3, 5 and 7 post-tibial fracture of wild-type mice and processed for single-nuclei RNAseq. **b.** UMAP projection of the integration of uninjured, day 1, day 3, day 5 and day 7 datasets. Eleven populations are identified and delimited by black dashed lines. **c.** Feature plot of the immune lineage score in the combined fracture datasets. **d**. Percentage of cells in the immune population per time point. **e.** UMAP projection of the subset of the immune cell clusters of the combined fracture datasets. The six populations are delimited by black dashed lines, and macrophage sub-populations are delimited by green dashed lines. **f.** Feature plots of the expression of *Arg1, Cd68, Mrc1* (CD206), and *Taco1* in the macrophage subpopulations**. g.** Percentage of cells in the immune cell populations per time point. uninj: uninjured. mac: macrophages. IIFCs: injury-induced fibrogenic cells. Osteob: osteoblasts.

### Macrophage dynamics during the inflammatory phase of bone healing

To further characterize the macrophage subpopulations in the early fracture healing environment, we performed flow cytometry and immunostaining analyses from uninjured periosteum and near adjacent muscle, and from the hematoma/callus at days 1, 3, 5 and 7 post-fracture from *Cx3cr1^GFP^*mice in which myeloid cells are marked by the GFP fluorescent reporter (Fig. 2a-g, Fig. S2). We confirmed the immune cell dynamics described in the snRNAseq analyses, with an increase of CD45+ and GFP+ cells at day 1 post-fracture, a peak at day 3 and a decrease at days 5 and 7 (Fig. 2b-f). In uninjured periosteum and muscle, the CD45+, GFP+, F4/80+ macrophage population was Arg1 and CD68 negative. At day 1, we identified a peak of Arg1^+^ macrophages and the majority of macrophages were Cd68^low^, corresponding to early macrophages (Fig. 2b-f, Fig. S3). At day 3, macrophages were CD68^high^ while the percentage of Arg1+ cells decreased, corresponding to intermediate macrophages. At day 5, macrophages were mostly CD68^low^ and we observed a continuous decrease of Arg1+ macrophages and a peak of CD206+ macrophages, corresponding to late macrophages 1 and 2. We performed immunostaining to determine the spatial distribution of these macrophage populations in the fracture callus. GFP+ cells localized in the entire callus/hematoma at days 1 and 3 post-fracture, and progressively decreased from day 5, specifically in the regions where cartilage islets and trabecular bone are forming (Fig. 2f). Interestingly, while Arg1+ and CD68+ macrophages were detected within the hematoma/callus, CD206+ macrophages were only detected in the muscle surrounding the fracture site, suggesting that this population is a muscle-specific macrophage subset (Fig. 2f-g). Overall, combined single nuclei transcriptomic, flow cytometry and histological analyses identified distinct macrophage subsets during the inflammatory phase of bone repair, with specific time and tissue patterning.

**Figure 2:**
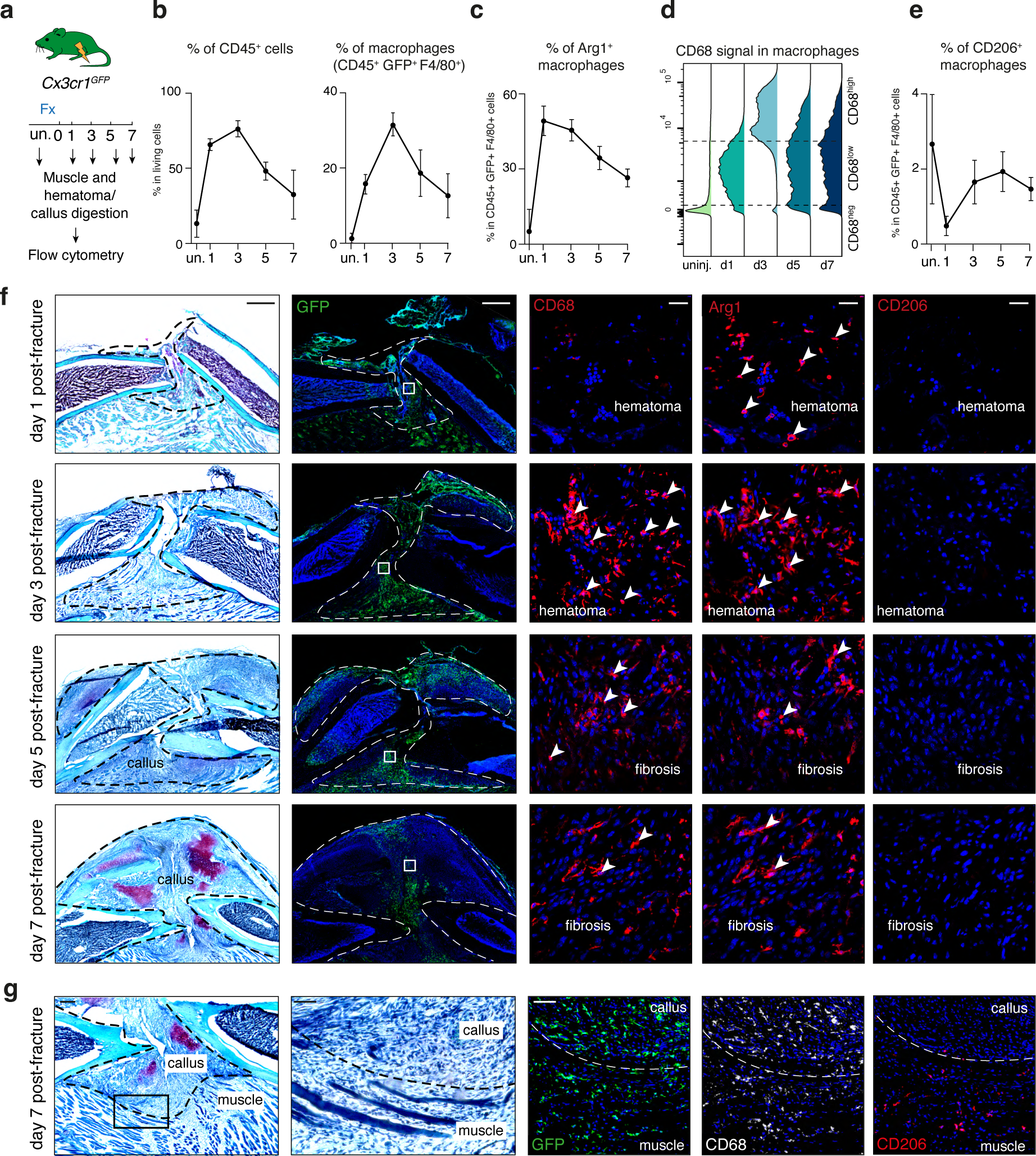
Dynamics of macrophage subsets during bone regeneration. **a.** Experimental design. Cells were isolated from uninjured muscle and periosteum and from the fracture environment (hematoma/callus, activated periosteum and skeletal muscle surrounding the fracture site) at 1, 3, 5 and 7 days post-fracture from *Cxc3cr1^GFP^* mice and analyzed by flow cytometry (n=3-7 per group). **b.** Percentage of CD45+ cells and macrophages (CD45+ GFP+ F4/80+ cells) in the uninjured periosteum/muscle and fracture environment at 1, 3, 5 and 7 days post-fracture of *Cx3cr1^GFP^* mice. **c.** Percentage of Arg1+ macrophages (CD45+ GFP+ F4/80+ Arg1+ cells). **d.** CD68 signal in the macrophage population (CD45+ GFP+ F4/80+ cells). **e.** Percentage of CD206+ macrophages (CD45+ GFP+ F4/80+ CD206+ cells). **f.** Left. Safranin’O staining and fluorescent images of GFP signal in longitudinal sections of tibial fracture at 1, 3, 5 and 7 days post-fracture. Middle and right: Immunofluorescence of CD68, Arg1 and CD206 on callus hematoma/fibrosis at 1, 3, 5 and 7 days post-fracture in *Cx3cr1^GFP^* mice showing cells labelled by Arg1 and/or CD68 (white arrowheads) (n=3 sections from 3 mice). **g.** Left: Safranin’O staining of tibial fracture section of *Cx3cr1^GFP^* mice at 7 days post-fracture. Right: Immunofluorescence of CD206 and CD68 on adjacent section show that CD206+ macrophages are only localized in the skeletal muscle surrounding the fracture site. Scale bars: f: high magnification: 1mm, low magnification: 25 μm. g. high magnification: 250 μm, low magnification: 100 μm.

### Distinct secretome and paracrine roles of the macrophage subsets during bone healing

Additional analyses of single nuclei transcriptomics defined the paracrine roles of macrophage subsets in bone healing. Connectome analysis showed that the paracrine role of immune cells is transient and mostly occurs at days 1 and 3 post-fracture (Fig. 3a). Among the immune cell populations, monocytes, early macrophages, intermediate macrophages, and late macrophages 1 had the highest interaction strength (Fig. 3b). We identified that macrophages express factors from several key signaling families, like interleukins, cytokines and PDGFs (Fig. S4a). Early macrophages expressed genes encoding pro-inflammatory factors including *Cxcl2, Ccl2, Ccl7, Ccl9 and Thbs1*, and predominantly at day 1 post-fracture (Fig. 3c)^29–31^. Intermediate macrophages expressed pro-repair factors such as *Tgfb1, Apoe,* and *Pf4* mostly at day 3 post-fracture (Fig. 3c)^31–34^. Finally, late macrophages 1 expressed anti-inflammatory factors like *Igf1, Gas6* and *Pdgfc*, mostly at day 5 post-injury (Fig. 3c)^30,35,36^. We observed the progressive shift from the pro-inflammatory phase at day 1 post-fracture to the anti-inflammatory phase at day 5 post-fracture. We also found that some pro-inflammatory genes like *Spp1* and *Nampt* were expressed by both early and intermediate macrophages, and some anti-inflammatory genes like *Il15* and *Ly86* were expressed by both intermediate and late macrophages, suggesting that intermediate macrophages represent the population at the interface between the pro- and anti-inflammatory phase of bone healing (Fig. S4b). Using CellChat package^37^, we identified the factors, including *Nampt, Spp1, Tgfb1*, *Pdgfc* and *Igf1*, expressed by early, intermediate, and late macrophages that can signal to SSPCs, IIFCs, chondrocytes and osteoblasts (Fig. 3d), unravelling multiple and dynamic interactions between macrophages and the SSPC, IIFC, chondrocyte and osteoblast populations.

**Figure 3:**
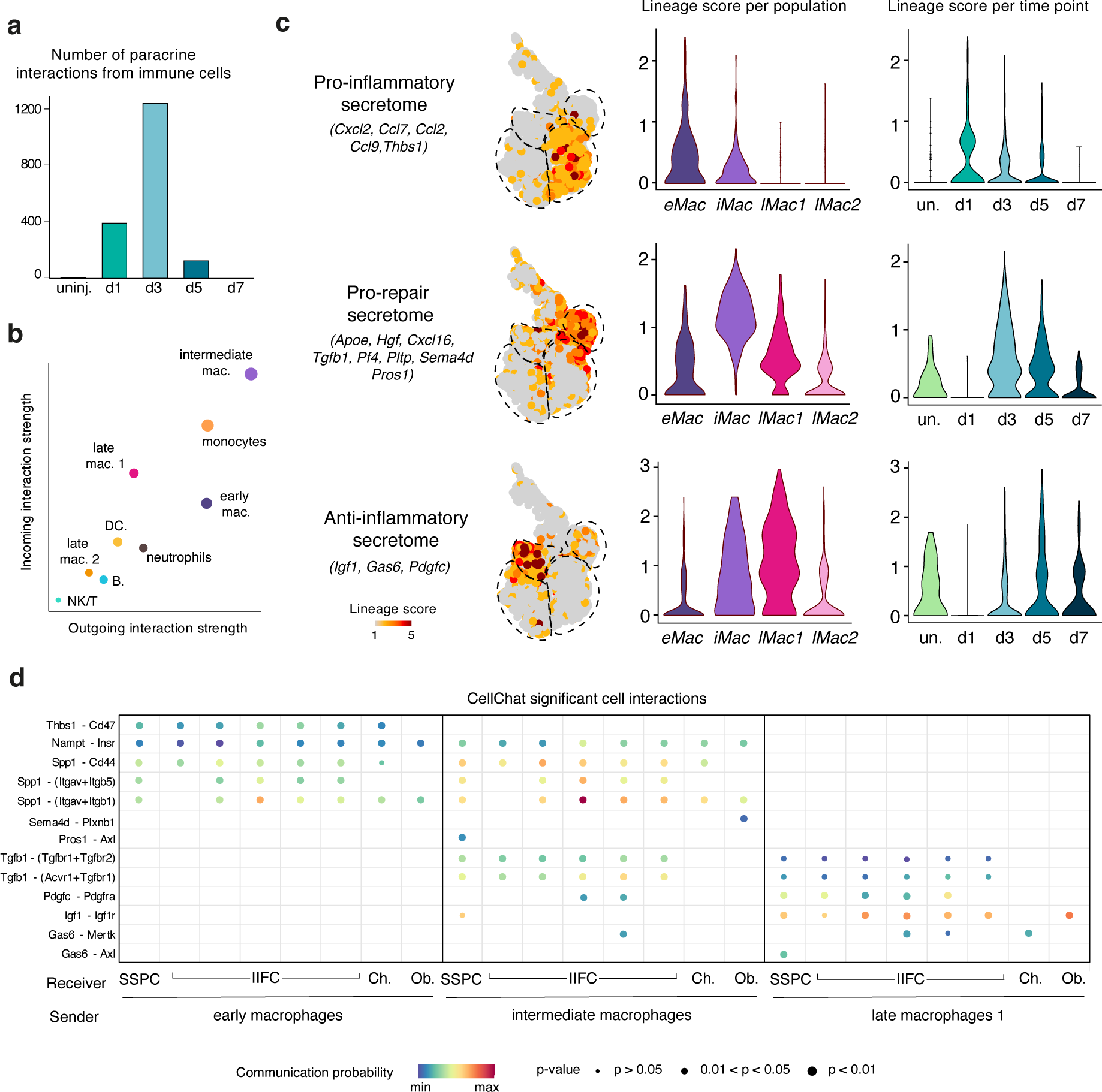
Distinct paracrine roles of macrophage subsets in the bone fracture environment. **a.** Number of paracrine interactions from immune cells per time point determined by Connectome package. **b.** Incoming and outgoing interaction plot of the immune cell populations showing that early macrophages, intermediate macrophages and late macrophages 1 are the main macrophage populations with a paracrine role after fracture. **c.** Feature plots (left) and violin plots per macrophage cluster (in all time points, middle) and per time point (in all macrophage clusters, right) of the score of the pro-inflammatory, pro repair and anti-inflammatory secretome in the subset of macrophages. **d.** Dot plot of the significant interactions from early, intermediate and late macrophages 1 to SSPCs, IIFCs, chondrocytes and osteoblasts determined by CellChat package. eMac: early macrophages, iMac: intermediate macrophages, lMac1: late macrophages 1, lMac2: late macrophages 2, un.: uninjured, Ch: chondrocytes, Ob.: osteoblasts.

### Fibrogenic and inflammatory response of SSPCs to bone fracture

Next, we focused our analyses on the fate of SSPCs during the inflammatory phase of bone repair. We performed a subset analysis of SSPCs, IIFCs, osteoblasts and chondrocytes (Fig. 4A). As previously reported, we observed that SSPCs differentiate in 2 consecutives steps by first engaging into an injury-induced fibrogenic state before undergoing either osteogenesis or chondrogenesis (Fig. 4a-b, Fig. S5)^4^. We observed that injury-induced fibrogenic cells (IIFCs) were divided in 2 subpopulations, one predominant at days 1 and 3 post-fracture (cluster 2) and one predominant at days 5 and 7 post fracture (clusters 3 to 5) (Fig. 4c-d). We performed gene regulatory network (GRN) analyses using SCENIC package (Single Cell rEgulatory Network Inference and Clustering) to identify regulons (transcription factors and their target genes) specifically active in the 2 IIFC subpopulations^38^. IIFCs from cluster 2 had active regulons linked to the stress response/cell activation (Junb, Fos), and to several pro-inflammatory signaling pathways including JAK-STAT (Stat1, Stat3), NFKB (Nfkb1, Rela) and Interferon regulatory factors (Irf1, Irf3). IIFCs from clusters 3 to 5 had active regulons linked to tissue regeneration, such as Six1, Meis1, Pax9 and to the resolution of inflammation, such as Pparg and Etv1. (Fig. 4e)^39–42^. These analyses revealed that IIFCs first exhibit a pro-inflammatory profile before switching to a pro-repair/anti-inflammatory profile. Cell interaction analyses showed that the secretome of IIFCs also changes during the steps of bone healing (Fig. S4c). Pro-inflammatory IIFCs (pIIFCs) expressed genes coding for pro-inflammatory factors including *Cxcl2, Cxcl5, Ccl2, Ccl5,* and *Csf1* while anti-inflammatory IIFCs (aIIFCs) expressed several pro-repair/anti-inflammatory genes such as *Igf1, Bdnf, Tgfb3, Gas6, Hgf* and *Inhba* (Fig. 4f). CellChat analysis established SSPCs and IIFCs as the main sources of signals for the immune cells after injury and more importantly for macrophages (Fig. S4d), indicating that IIFC signaling can also modulate the inflammatory response to bone fracture. Using our previously published scRNAseq datasets^8^, we confirmed that muscle SSPCs also adopt a pIIFC profile followed by an aIIFC profile, associated with secretion of pro-inflammatory and anti-inflammatory factors respectively (Fig. S6a-d). Overall, during their fibrogenic response to bone fracture SSPCs adopt successive inflammatory profiles that parallel the temporal dynamic of macrophages.

**Figure 4:**
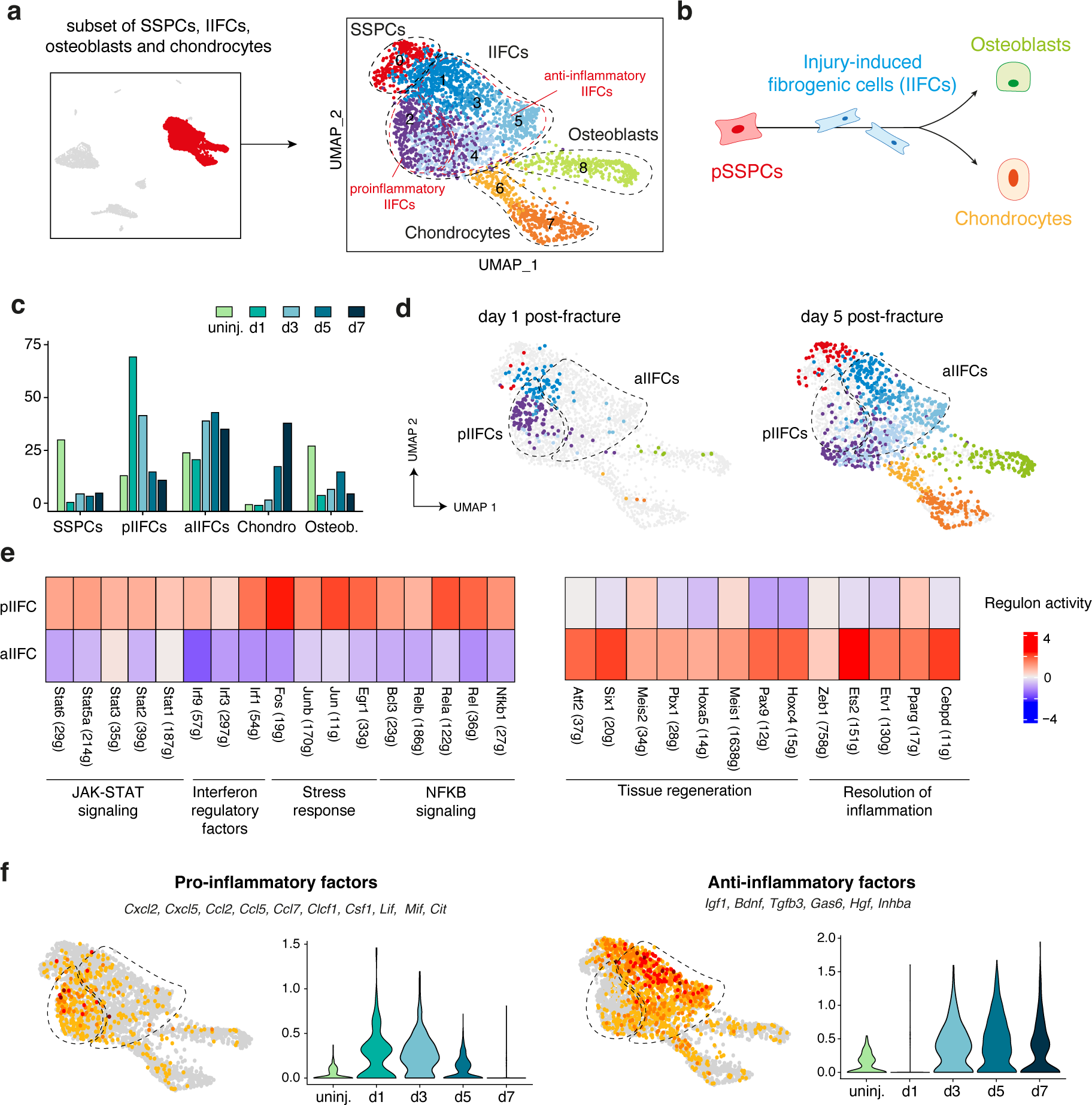
Inflammatory profile of SSPCs in response to bone fracture. **a.** UMAP projection of the subset of SSPCs, IIFCs, osteoblasts and chondrocytes from the combined fracture dataset. **b.** Schematic representation of the differentiation trajectory of pSSPCs after bone fracture. **c.** Percentage of cells in the different populations per time point. **d.** UMAP projection of the cells from the day 1 and day 5 post-fracture datasets in the integrated dataset. **e.** Heatmap of the regulon activity upregulated in pro-inflammatory IIFCs (pIIFCs, left) and upregulated in anti-inflammatory IIFCs (aIIFCs, right). **f.** Feature plots and violin plots per time point of the score of the expression of pro-inflammatory factors (left) and anti-inflammatory factors (right). pSSPCs: periosteal skeletal stem progenitor cells.

### Prolonged inflammatory response in musculoskeletal trauma

We previously showed that musculoskeletal traumatic injury (i.e. combined fracture and adjacent muscle injury) causes fibrotic accumulation in the fracture callus and subsequent bone non-union (Fig. 5a)^8^. At 21 days post-fracture, the callus was fully ossified with almost complete resorption of fibrous tissue^8^ and we did not detect CD68+ and CD206+ macrophages in callus fibrosis. However, GFP+ cells were detected in the newly formed bone marrow tissue surrounding bone trabeculae (Fig. 5a-c). After MTI, we observed accumulation of GFP+ macrophages expressing CD68 and CD206 in the persistent fibrous tissue within the callus (Fig. 5b-c). This suggested that the macrophage dynamic is perturbed in MTI and could contribute to fibrotic accumulation. To investigate the impact of MTI on the immune cell response to injury, we performed flow cytometry analysis of the cells from fracture hematoma/callus and surrounding muscles after MTI and compared with the fracture data from Fig. 1 (Fig. 5d). At day 1 post-injury, we observed similar percentages of CD45+ cells in MTI and fracture, but decreased percentages of macrophages, Arg1+ macrophages and CD68^low^ macrophages in MTI, indicating a delay in macrophage recruitment or activation after MTI (Fig. 5e-f). At day 3 post-injury, no differences were observed between fracture and MTI. At day 5 and 7, we detected a decrease in immune cells in the fracture callus compared to day 3, while in MTI the percentage of CD45+, GFP+ cells, total macrophages, Arg1+, and CD68^high^ macrophages remained high and was significatively increased compared to fracture (Fig. 5e-f, Fig. S2c). In addition, the percentage of CD206+ macrophages continued to increase and remained high in MTI compared to fracture. Analysis on tissue sections confirmed the accumulation of macrophages in the callus and skeletal muscle surrounding the fracture site (Fig. 5g), showing an altered immune response marked by a delay in the resolution of inflammation at days 5 and 7 post-MTI.

**Figure 5:**
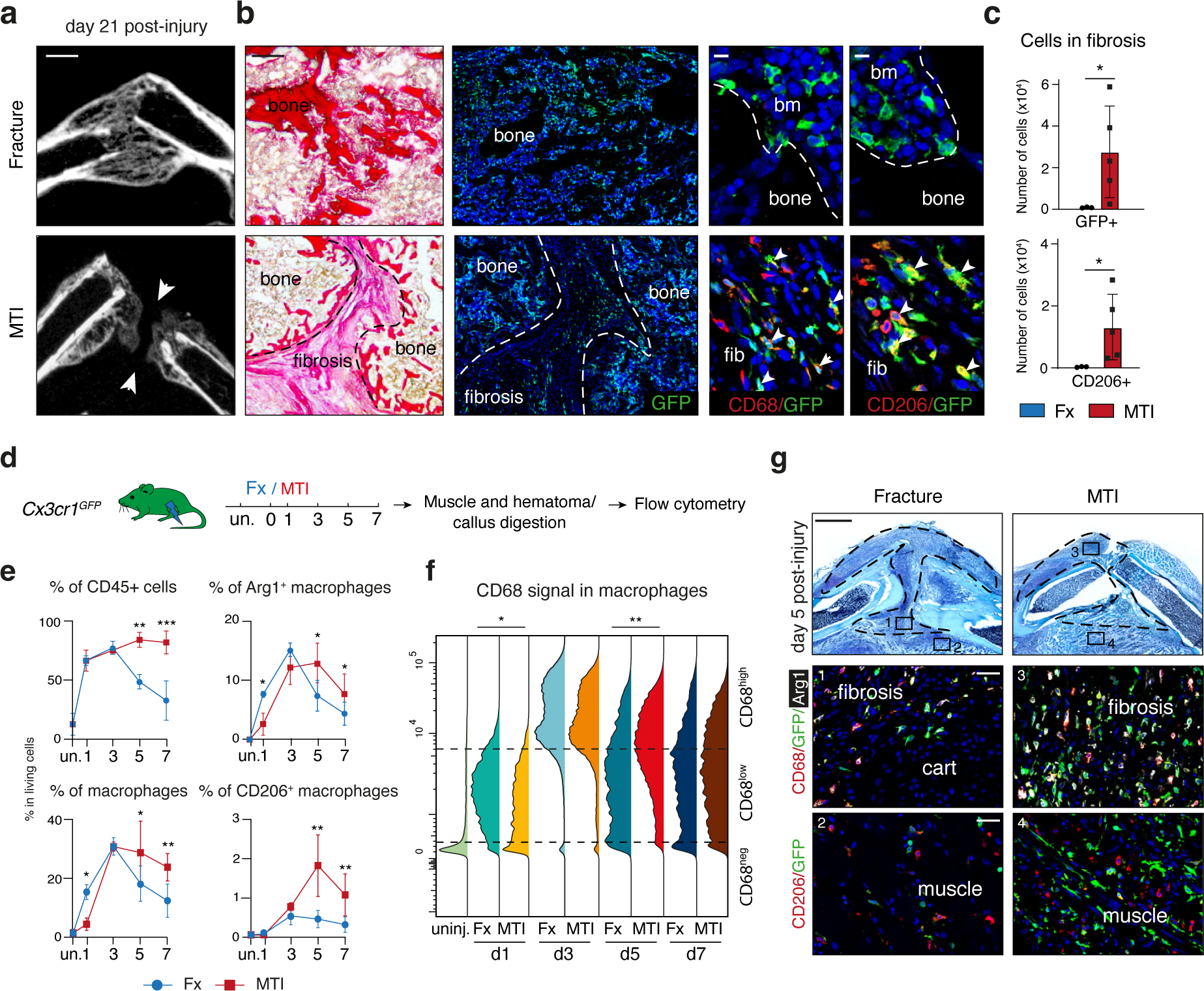
Musculoskeletal trauma alters macrophage dynamic during bone repair. **a.** Left: microCT images of day 21 post-fracture (top) or post-MTI (bottom) callus showing absence of bone union in MTI (white arrows). **b.** Low magnification of picrosirius staining and GFP signal on adjacent section of 21-days post-injury callus of *Cx3cr1^GFP^*mice and high magnification of the callus showing GFP+ CD206+ and GFP+ CD68+ cells in the fibrosis after MTI (white arrows). **c**. Number of GFP+ and CD206+ cells in the callus fibrosis at day 21 post-fracture or MTI (n=3-4 per group). **d.** Experimental design. Cells were isolated from muscles and hematoma/callus at day 1, 3, 5 and 7 post-fracture or MTI of *Cx3cr1^GFP^* mice and analyzed by flow cytometry (n=3-8 per group). **e.** Percentage of CD45+ cells, macrophages (CD45+ GFP+ F4/80+), and percentage of Arg1+ and CD206+ macrophages in uninjured tissue and at 1, 3, 5 and 7 days post-fracture or MTI of *Cx3cr1^GFP^* mice. **f.** CD68 signal in the macrophage population (CD45+ GFP+ F4/80+ cells). Statistical difference was determined between the median of fluorescent signal per sample. **g.** Top: Safranin’O staining of day 5 post-fracture or MTI callus. Middle-bottom: Low magnification of the callus section of *Cx3cr1^GFP^*mice and high magnification of the callus and muscle showing increased amount of GFP+ CD206+ and GFP+ CD68+ cells in the fibrosis after MTI. p-value: *: p<0.05, **; p<0.01.bm: bone marrow, fib: fibrosis. Scale bars: a. 1mm. b. low magnification: 1mm, high magnification: 10 μm. g. low magnification: 1mm, high magnification: 100 μm.

To elucidate the altered inflammatory response in MTI, we focused our analyses on the day 5 post-injury by generating a snRNAseq dataset of the fracture callus 5 days post-MTI (Fig. 6a). We integrated with the day 5 post-fracture dataset and identified the same cell populations as in the bone repair atlas (Fig. 6a). The percentages of IIFCs, chondrocytes and osteoblasts were reduced in the MTI dataset compared to fracture, while the percentage of immune cells was increased (Fig. 6b-c), confirming the persistent immune response at day 5 in MTI. Detailed analysis of the immune cells showed an overall increase of immune cells, but specifically pro-inflammatory early macrophages (Fig. 6d-e). Gene ontology (GO) analyses indicated enrichment in functions related to macrophage activation and proinflammatory activity (Fig. 6f). More precisely, while eMac in fracture showed upregulation of GO related to the regulation of cellular processes, eMac in MTI showed upregulation of GO linked to macrophage function, revealing their increased activity (Fig. S7). We observed an increased expression of pro-inflammatory signals by immune cells and a decrease in pro-repair and anti-inflammatory factors expression (Fig. 6g). Therefore, MTI alters the inflammatory response to fracture, with a delay in macrophage recruitment, a prolonged pro-inflammatory phase and a delayed anti-inflammatory phase and resolution of inflammation.

**Figure 6:**
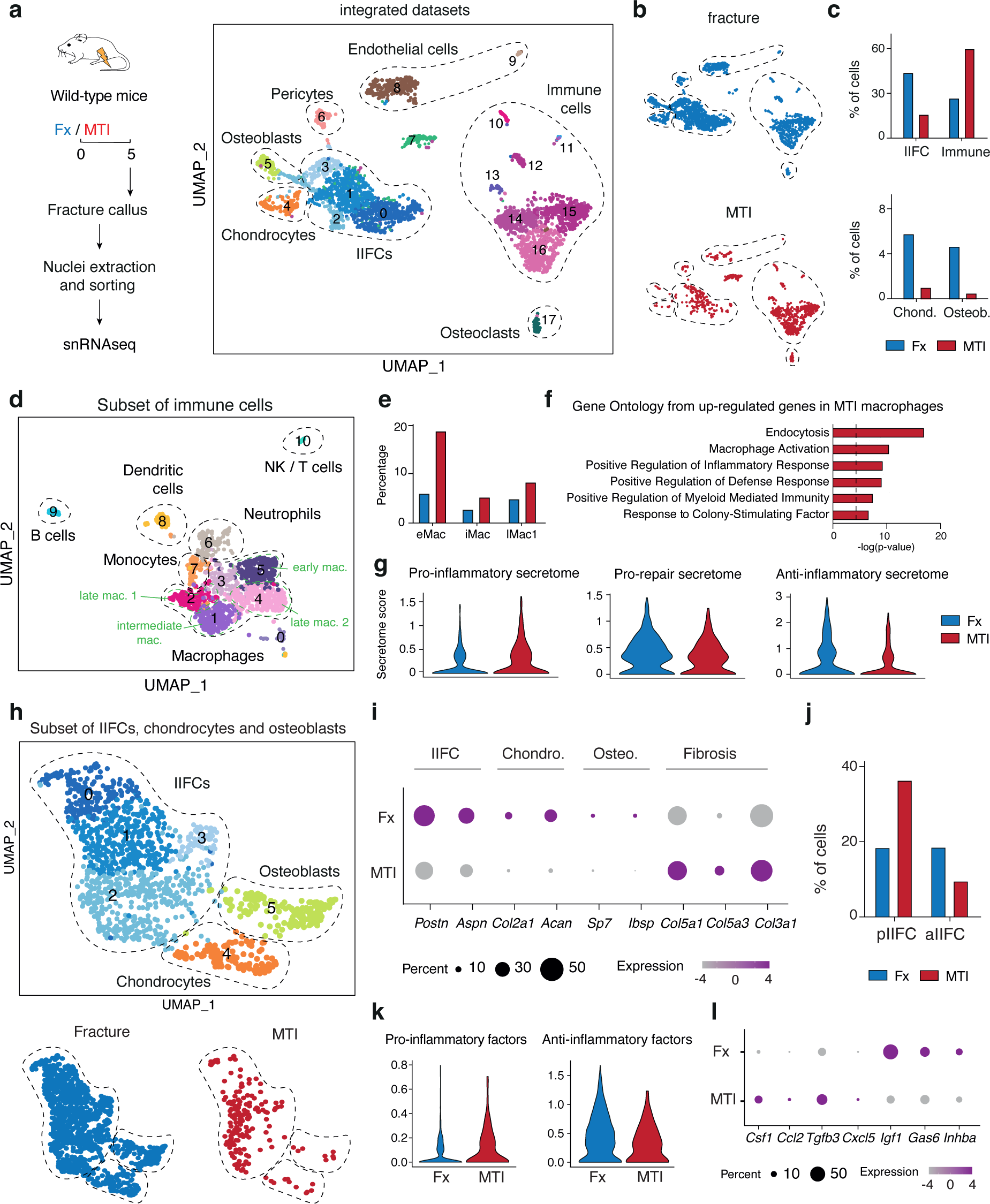
Prolonged pro-inflammatory state of macrophages and SSPCs after musculoskeletal trauma. **a.** Experimental design. Nuclei were extracted from the fracture callus at day 5 post-fracture or post-MTI and processed for snRNAseq. UMAP projection of the integration of the day 5 post-fracture and post-MTI datasets. **b.** UMAP projection of the integrated dataset separated by dataset of origin. **c.** Percentage of cells in the IIFC, immune cell, chondrocyte and osteoblast population per dataset. **d.** UMAP projection of subset of immune cells from the day 5 post-fracture and MTI datasets **e.** Percentage of early, intermediate and late macrophages in the total dataset from day 5 post-fracture and MTI. **f.** Gene ontology (GO) of the upregulated genes in macrophages (clusters 1 to 5) from MTI dataset compared to fracture dataset. **g.** Violin plots of the score of pro-inflammatory, pro-repair and anti-inflammatory secretome in the subset of immune cells. **h.** Top: UMAP projection of the subset of IIFCs, chondrocytes, and osteoblasts from the day 5 post-fracture and post-MTI datasets. Bottom: UMAP projection per dataset of origin**. i.** Dot plot of genes markers in IIFCs, chondrocytes, osteoblasts and fibrosis genes specifically upregulated in MTI**. j.** Percentage of pro-inflammatory IIFCs (pIIFCs) and anti-inflammatory IIFCs (aIIFCs) in the fracture and MTI dataset. **k.** Violin plots of the expression of pro-inflammatory and anti-inflammatory factors in the subset dataset. **l.** Dot plot of genes encoding for pro-inflammatory factors and upregulated in MTI and genes encoding for anti-inflammatory factors and upregulated in fracture. pIIFC: pro-inflammatory injury induced fibrogenic cells, aIIFC: anti-inflammatory injury induced fibrogenic cells, eMac: early macrophages, iMac: intermediate macrophages, lMac1: late macrophages 1. Chond.: chondrocyte. Osteob.: Osteoblast.

### Altered immune response of SSPCs in musculoskeletal trauma

We next focused the snRNAseq analyses on the impact of MTI on SSPC differentiation. We generated a subset of the IIFCs, chondrocytes and osteoblasts (Fig. 6h). We observed a strong reduction of IIFCs, chondrocytes and osteoblasts marker gene expression in MTI compared to fracture, and identified upregulated ECM genes in MTI (*Col5a1, Col5a3, Col3a1*) (Fig. 6i). Specifically, the percentage of pro-inflammatory IIFCs (pIIFCs) was increased in MTI and the percentage of anti-inflammatory IIFCs (aIIFCs) was decreased, indicating a delay in the switch from pIIFCs to aIIFCs (Fig. 6j). In parallel, we observed increased expression of pro-inflammatory factors, such as *Csf1*, *Ccl2* and *Cxcl5,* and decreased expression of anti-inflammatory factors (*Igf1, Gas6, Inhba*) (Fig. 6k-l). We confirmed these observations using scRNAseq datasets of SSPCs in uninjured muscles and muscles adjacent to the fracture site at days 3 and 5 post-fracture or MTI (Fig. S6e). We also identified pIIFC and aIIFC populations at day 3 post-injury with a delay in the switch from pIIFCs to aIIFCs at day 5 post-injury in the MTI dataset, correlated with increased expression of pro-inflammatory factors and decreased expression of anti-inflammatory factors (Fig. S6f-h). Overall, MTI causes an altered immune response at the fracture site, with delayed macrophage recruitment, delayed resolution of inflammation in SSPCs and macrophages, and accumulation of macrophages in the fibrotic tissue at late stages of repair.

### Macrophages promote fibrotic accumulation in musculoskeletal trauma

We then assessed functionally the contribution of macrophages to bone fibrosis and nonunion in MTI. We first assessed bone healing in *Ccr2* knock-out mice (*Ccr2*^-/-^) where systemic recruitment of macrophages is impaired^43^. We induced MTI in *Ccr2*^-/-^ mice and *Ccr2*^+/+^ controls and observed 75% of bone union in *Ccr2^-/-^* mice compared to 20% in controls 35 days post-injury, suggesting that reducing macrophage recruitment in MTI can ameliorate bone healing (Fig. 7a-b). Improved healing in *Ccr2^-/-^*mice was associated with a drastic reduction in callus fibrosis 21 days post-fracture and fewer CD206+ and CD68+ cells in the callus (Fig. 7c-d). We then used an inducible approach to deplete macrophages from day 2 to day 4 post-MTI to prevent the prolonged inflammation observed in MTI. We used *LysM^Cre^; R26^tdTom^* mice that efficiently mark all macrophage populations including CD68+ and CD206+ cells accumulating in the fibrotic tissue of MTI (Fig. S8a). Diphteria toxin injection in *LysM^Cre^; R26^tdTom/IDTR^* mice from days 2 to 4 post-MTI decreased the percentage of macrophages (F4/80+ CD45+ Tomato+ cells) and the percentage of Arg1+ and CD68^High^ macrophages at day 5 post-fracture (Fig. S8b). By day 35, we observed improved bone healing, with bone union/semi-union observed in 50% of *LysM^Cre^; R26^tdTom/IDTR^* mice compared to 0% in *LysM^Cre^; R26^tdTom^*control mice (Fig. 7e-f). Macrophage depletion led to significant decrease in fibrosis volume and fewer CD206+ and CD68+ macrophages in the callus fibrosis of depleted mice compared to control mice at 21 days post-MTI (Fig. 7g-h).

**Figure 7:**
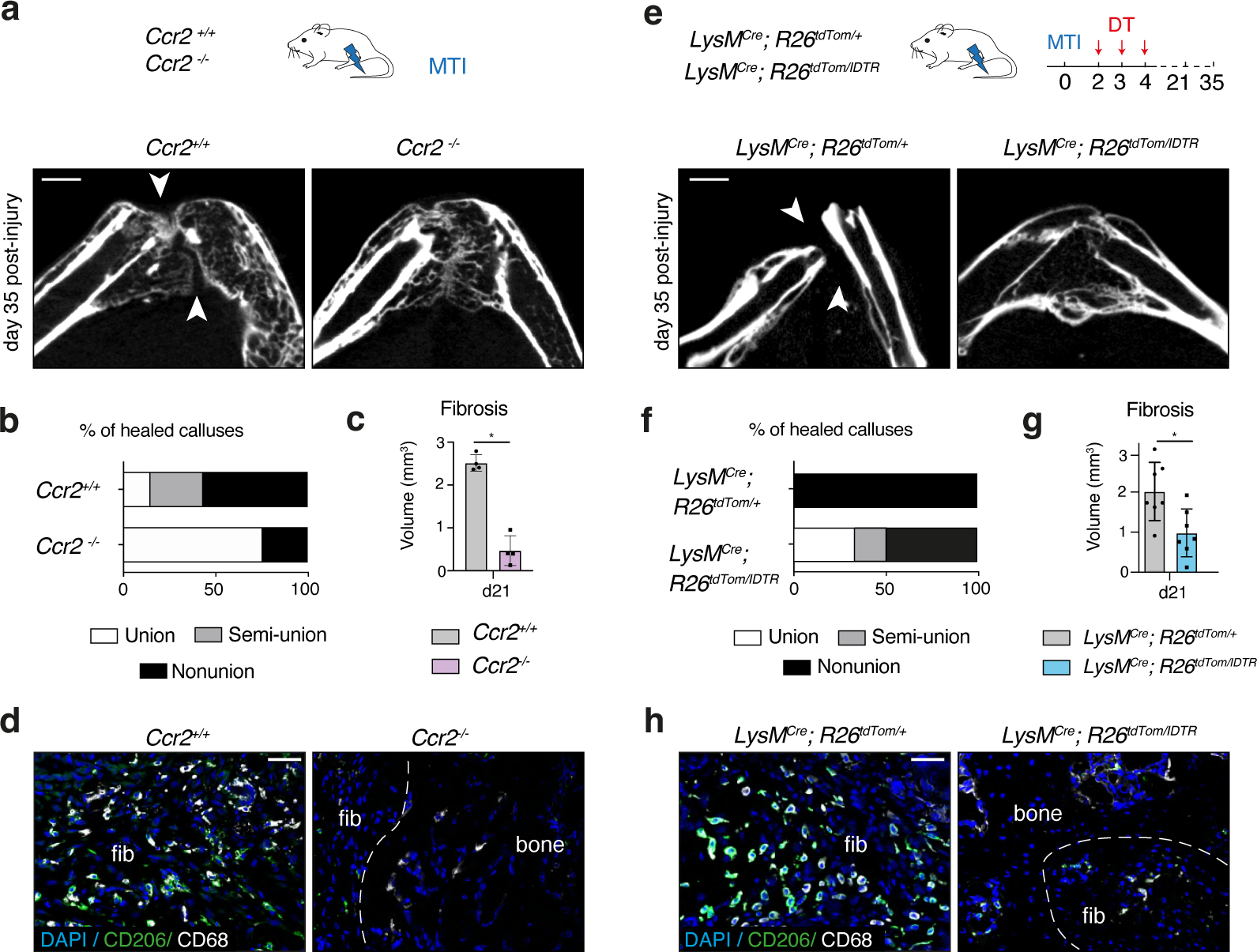
Macrophage depletion prevents fibrotic accumulation in musculoskeletal trauma. **a.** Experimental design. MTI was induced in *Ccr2*^+/+^ and *Ccr2*^-/-^ mice, in which macrophages fail to infiltrate tissues during inflammation. Representative microCT scan of 35 days post-MTI callus from *Ccr2*^+/+^ and *Ccr2*^-/-^ mice, showing nonunion in *Ccr2*^+/+^ mice (white arrows) and bone bridging in *Ccr2*^-/-^ mice. **b.** Percentage of healed calluses from *Ccr2*^+/+^ and *Ccr2*^-/-^ mice showing bone union (white), semi-union (grey) or nonunion (black) on microCT scan at day 35 post-MTI (n=4-5 mice per group). **c.** Volume of fibrosis in the callus of *Ccr2*^+/+^ and *Ccr2*^-/-^ mice at 21 days post-MTI. **d.** Immunofluorescence on callus sections of *Ccr2*^+/+^ and *Ccr2*^-/-^ mice at 21 days post-MTI showing presence of CD206+ and CD68+ macrophages only in the fibrosis of *Ccr2^+/+^* mice (white arrows). **e.** Experimental design. MTI was induced to *LysM^Cre^; R26^tdTom^* and *LysM^Cre^; R26^tdTom/IDTR^* mice, and diphteria toxin (DT) was injected at days 2, 3 and 4 post-injury to induce macrophage depletion. Representative microCT scan of 35 days post-MTI callus from *LysM^Cre^; R26^tdTom^*and *LysM^Cre^; R26^tdTom/IDTR^* mice, showing nonunion in *LysM^Cre^; R26^tdTom^*mice (white arrows) and bone bridging in *LysM^Cre^; R26^tdTom/IDTR^*mice. **f.** Percentage of healed calluses from *LysM^Cre^; R26^tdTom^* and *LysM^Cre^; R26^tdTom/IDTR^* mice showing bone union (white), semi-union (grey) or nonunion (black) on microCT scan at day 35 post-MTI (n=7 mice per group). **g.** Volume of fibrosis in the callus of *LysM^Cre^; R26^tdTom^* and *LysM^Cre^; R26^tdTom/IDTR^* mice at 21 days post-MTI. **h.** Immunofluorescence on callus section of *LysM^Cre^; R26^tdTom^* and *LysM^Cre^; R26^tdTom/IDTR^* mice at 21 days post-MTI showing presence of CD206+ and CD68+ macrophages only in *LysM^Cre^; R26^tdTom^* mice (white arrows). Scale bars: a-e: 1mm, d-h: 100 μm. p-value: *: p<0.05.

### Pharmacological CSF1R inhibition improves bone healing in musculoskeletal trauma

Given the direct involvement of macrophages in the MTI phenotype, we sought to test a pharmacological approach by targeting macrophages to improve MTI outcome. The FDA-approved CSF1R inhibitor Pexidartinib was previously shown to efficiently deplete Tumor-associated macrophages (TAMs) in osteosarcomas^44^. In fracture healing, *Csf1r* expression is limited to macrophages and osteoclasts and increased in MTI (Fig. S8c-f). Further, *Csf1* was upregulated in IIFCs in MTI suggesting that inhibiting CSFR1 may be an efficient strategy to decrease macrophage function in MTI (Fig. 6l). We induced MTI in wild type mice and treated them daily with oral gavage of Pexidartinib or vehicle from days 3 to 14 post-injury (Fig. 8a). At 21 days post-fracture, we observed improved bone healing with a significant increase in the number of bridged sides in Pexidartinib-treated compared to vehicle-treated mice (Fig. 8b-c). CSF1R inhibition led to increased mineralized bone and cartilage in the callus and reduced fibrosis accumulation compared to control (Fig. 8d-e). In addition, we observed fewer CD206+ and CD68+ cells in the callus of Pexidartinib treated mice (Fig. 8f). This shows that pharmacological CSF1R inhibition is a promising strategy to improve bone healing after musculoskeletal trauma.

**Figure 8:**
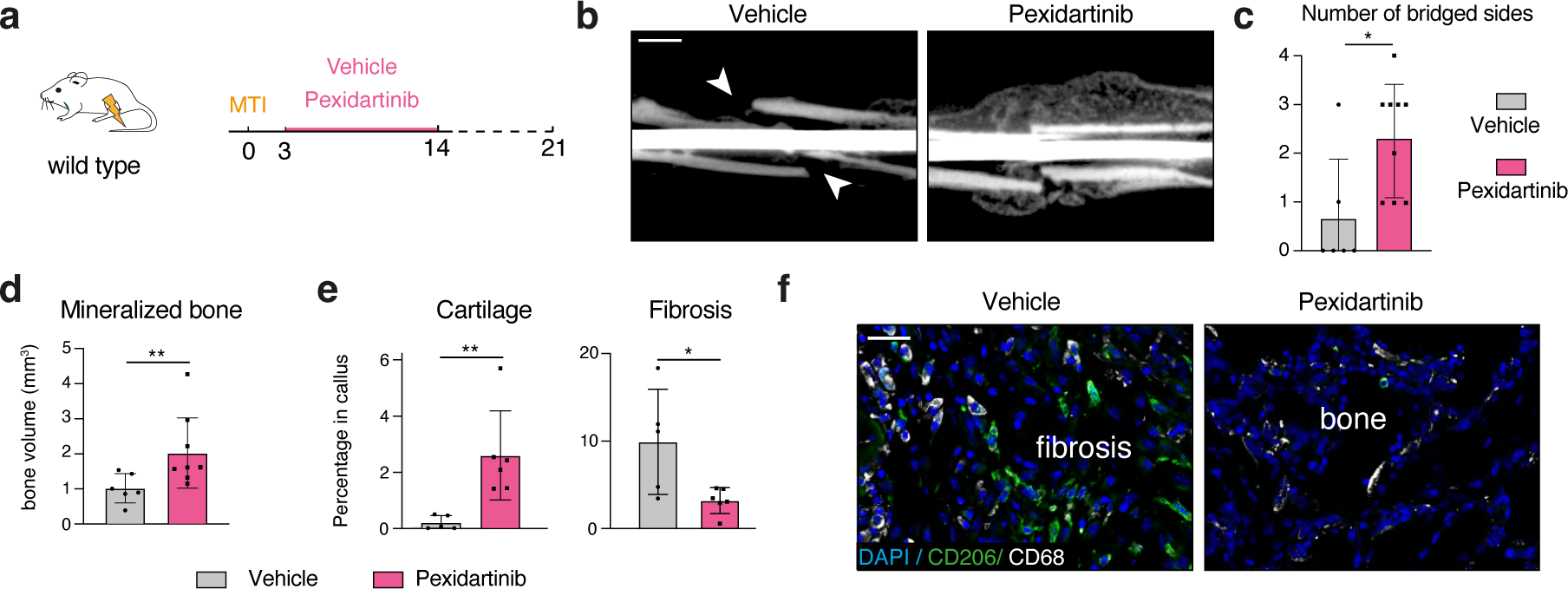
CFS1R inhibition prevents fibrous nonunion after musculoskeletal trauma. **a.** Experimental design. Wild-type mice were treated by oral gavage of Pexidartinib or vehicle from days 3 to 14 post-MTI. **b.** Representative microCT scan of 21 days post-MTI callus from mice treated with Pexidartinib or vehicle, showing nonunion in vehicle-treated mice (white arrows) and bone bridging in Pexidartinib-treated mice (n=6-9 mice per group). **c.** Number of bridged sides of day 21-post MTI callus of mice treated with Pexidartinib or vehicle. **d.** Volume of mineralized bone in 21 days post-MTI callus of mice treated with Pexidartinib or vehicle (n=6-9 mice per group). **e.** Percentage of cartilage and fibrosis in day 21-post MTI callus of mice treated with Pexidartinib or vehicle. **f.** Immunofluorescence on the callus section of vehicle- and Pexidartinib-treated mice at 21 days post-MTI showing presence of CD206+ and CD68+ macrophages only in vehicle-treated mice (white arrows). Scale bars: b: 1mm, e: 100 μm. p-value: *: p<0.05, **<0.01.

## Discussion

Bone regeneration depends on an efficient differentiation of SSPCs, coordinated by multiple cellular interactions in the fracture environment. SSPCs exhibit plasticity in cell fate commitment during the early phase of bone regeneration, which coincides with the inflammatory phase of healing. Similarly, immune cells possess incredible capacities to change their cellular phenotypes to adapt and respond to tissue damage. Yet the immune cell dynamics and interactions with SSPCs during fracture repair is not fully understood. Here, we combined single-nuclei transcriptomics, flow cytometry and in vivo analyses to characterize the rapid changes in the immune cell populations during the inflammatory response to bone injury. We described that immune cells from the myeloid and lymphoid lineages appear from day 1 post-injury, peak at day 3 before regressing from day 5 after fracture. Macrophages were the predominant immune population and switched from a pro-inflammatory to an anti-inflammatory profile as previously reported^20,45^. However, our transcriptomic analyses revealed a greater macrophage heterogeneity than previously described. We defined four major successive macrophage subtypes expressing both classical and alternatively activated macrophage markers, such as Arginase 1, CD206 and CD68. Although Arginase 1 was considered as an anti-inflammatory macrophage marker, we and others found that *Arg1* is expressed by early pro-inflammatory macrophages during tissue repair^45,46^. Further analyses will aim to decipher if the macrophage subtypes identified are different cell types or cells undergoing progressive transitions. Each macrophage subtype was associated with a distinct secretome, and interaction analyses identified several signals from macrophages to SSPCs, IIFCs, osteoblasts and chondrocytes such as SPP1, TGFβ1, IGF1, GAS and PDGFc. While some of these factors were previously shown to modulate bone repair^47,48^, the role of others still requires further analyses.

Remarkably, we observed pro- and anti-inflammatory phases during the fibrogenic response of SSPC, that correlate with the immune cell response. Our study shows that SSPCs from the periosteum and the skeletal muscle can activate immune-related signaling (NFKB, JAK-STAT) during the fibrogenic phase in IIFCs prior to osteochondrogenic differentiation. This transitional stage, where both the fibrogenic response and the immune response coincide, is crucial for SSPC cell fate decision, revealing the direct impact of immune regulation on SSPC differentiation. In addition, IIFCs secrete pro- and anti-inflammatory factors, including cytokines and interleukins. While bone marrow SSPCs are known to be able to modulate inflammation^49,50^, further analyses will aim to understand to which extent IIFCs derived from periosteum and muscle are able to regulate the inflammatory phase of bone repair and modulate macrophage activation and identity. Similar phenomenon was recently described in wound healing and oral chronic disease, highlighting fibroblast regulation of inflammation as a key factor in tissue homeostasis and repair^46,51–53^. The parallel inflammatory response in SSPCs and macrophages demonstrate the critical crosstalk between these 2 populations at the early steps of bone healing and place macrophages as a central player in bone regeneration.

These critical functional interactions between SSPCs and macrophages are further exemplified by the analyses of the fibrous nonunion phenotype associated with musculoskeletal traumatic injury. In the mouse MTI model, we uncovered an altered immune response with prolonged pro-inflammatory phase and delayed anti-inflammatory phase. Both macrophage and SSPC dynamics were delayed leading to sustained expression of genes encoding for pro-inflammatory factors and reduced expression of anti-inflammatory genes. The altered early inflammatory phase led to accumulation of CD206+ macrophages in callus fibrosis at late stages of healing. Interestingly, CD206+ cells, corresponding to the late macrophage 1 population, are only found in the skeletal muscle during normal bone healing. The pro-fibrotic role of this CD206+ macrophage population in MTI correlates with previous reports showing the deleterious effect of muscle CD206+ macrophages on muscle regeneration^54^. Immune response dysregulation was also described in delayed bone repair associated with aging or skeletal fluorosis^55,56^, confirming the importance of a balanced initiation and resolution of the inflammatory phase during bone repair for successful healing.

The pivotal role of macrophages in driving fibrosis accumulation suggests that modulating their functions can prevent fibrous nonunion. Genetic depletion of macrophages confirmed macrophages as an optimal target to improve healing outcome in musculoskeletal trauma and targeting specific macrophage subtypes or mediators may be of particular interest for clinical applications. Our findings provide a proof of concept that pharmacological inhibition of macrophages using the CSF1R inhibitor Pexidartinib can prevent fibrous nonunion. Pexidartinib was previously shown to reduce macrophage recruitment leading to reduced tumorigenesis and fibrosis accumulation in lung and heart injury^28,57,58^. Indeed, Pexidartinib treatment successfully led to reduced fibrosis accumulation along with improved healing in our mouse preclinical model, opening exciting new perspectives for the treatment of musculoskeletal trauma.

## Materials and Methods

### Mice

C57BL/6ScNj, B6.129P-CX3CR1^tm1Litt^/J (*Cx3cr1*^gfp^)^59^; B6.129P2-Lyz2^tm1(cre)Ifo^/J (*LysM^Cre^*)^60^*; R26tdTomato* (*R26^tdTom^*)^61^, C57BL/6-Gt(ROSA)26Sor^tm1(HBEGF)Awai^/J (*R26^IDTR^*)^62^*, B6.129S4-CCR2^tm1^ ^Ifc^*/J (*CCR2^-/^*^-^)^43^ were obtained from Jackson Laboratory (Bar Harbor, ME). Mice were kept in a pathogen-controlled environment in animal facilities at IMRB, Creteil, Paris and Karolinska Institute, Stockholm, Sweden. All procedures performed were approved by Paris Est Creteil University and Karolinska institute ethical Committees. Males and females were mixed in experimental groups. Bone injury and tissue collection for graft and digestion were performed on 10-to 14-week-old mice.

### Tibial fracture and Musculoskeletal Traumatic Injury (MTI)

Mice were anesthetized with an intraperitoneal injection of Ketamine (50 mg/mL) and Medetomidine (1 mg/kg) and received a subcutaneous injection of Buprenorphine (0.1 mg/kg) for analgesia. Open non-stabilized tibial fractures were performed as described. The right hindlimb was shaved, sanitized and the skin was incised to expose the tibia. An osteotomy was performed in the mid-diaphysis by cutting the bone. For MTI, the skin was incised and skeletal muscles surrounding the tibia were separated from the bone. The muscles, including tibialis anterior (TA), tibialis posterior, extensor digitorum longus (EDL), soleus, plantaris and gastrocnemius muscles, were subjected to mechanical injury by compression for 5 to 10 seconds along their entire length using a hemostat in a standardized and reproducible procedure^8^. The wound was sutured using non-resorbable sutures. An intraperitoneal injection of atipamezole (1 mg/kg) was performed to revive the mice and two additional analgesic injections were performed in the 24 hours following surgery.

### Diphteria toxin induced cell depletion

To genetically deplete myeloid cells, we used *LysM^Cre^; R26^tdTom/iDTR^* mice, in which LysM-expressing myeloid cells also express the diphteria toxin receptor (DTR). *LysM^Cre^; R26^tdTom/iDTR^* mice received three intraperitoneal injections with diphtheria toxin (DT, 20 ng/ml) (Sigma, St. Louis, MO) at 48 h, 72 h and 96h post MTI, inducing cell death in cells expressing the DTR (i.e. myeloid cells in our model). Control littermates (*LysM^Cre^; R26^tdTom/tdTom^*) received the same regimen.

### Pexidartinib treatment

After MTI, mice were orally dosed by gavage and received pexidartinib (PLX3397, Biorbyt, UK) at a dose of 50 mg/kg/day, or vehicle (0.5% HPMC, 1% Tween80 and 5% DMSO) starting from the third day post MTI and for 12 days.

### MicroCT and bone union scoring

Callus samples were scanned at the Small Animal Platform of Paris Cité University (EA2496, Paris Cité) using X-ray micro-CT device (Quantum FXCaliper, Life Sciences, Perkin Elmer, Waltham, MA) with the X-ray source at 90 kV and 160 μA. A 10 mm field of view and 20 μm voxel size was used. Horos software was used to visualize the callus and evaluate the number of bridged sides in sagittal and longitudinal axes. Bone union corresponds to a callus bridged in 3 or 4 sides, semi-union to 2 sides, and nonunion to 1 or 0 side ^8^. Mineralized bone volume in the total callus was measured using CTan software.

### Tissue sample processing and histology

Samples were processed as previously described^63^. Fractured tibias were collected and fixed in ice-cold 4% paraformaldehyde 24 hours, decalcified 2 weeks in 19% EDTA (EU00084, Euromedex), placed 24h in 30% sucrose (200-301-B, Euromedex) gradient and embedded in cryoprotectant. Samples were cryosectionned throughout the entire callus in 10µm thick sections. All thirtieth sections were defrosted, rehydrated and stained with Picrosirius, Masson’s Trichrome or Safranin’O staining as described below. After staining, sections were dehydrated with successive ethanol and NeoClear incubations. Slides were mounted using NeoMount medium (1.09016.0100, VWR, USA) and pictured using a Zeiss Imager D1 AX10 light microscope (Carl Zeiss Microscopy GmbH).

### Picrosirius staining (PS)

Sections were stained in PicroSirius solution, 0.1% of Direct Red 80 (43665-25G, Merck) in picric acid (80456, Merck) for 2 hours.

### Safranin’O staining (SO)

Sections were stained with Weigert’s solution for 5 min, washed in tap water for 3 min and stained with 0.02% Fast Green for 30 seconds (F7252, Merck), followed by 1% acetic acid for 30 seconds and Safranin’O solution for 45 min (S2255, Merck).

### Histomorphometric analysis

Histomorphometric analysis was performed throughout the entire callus as described previously ^7^. The cartilage and fibrosis surfaces were determined every thirtieth sections using ZEN software v1.1.2.0 (Carl Zeiss Microscopy GmbH) on SO and PS staining respectively. Volumes were calculated using the formula: 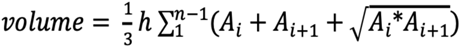 with A_i_ and A_i+1_ being the areas of cartilage and fibrosis in following sections, h was the distance between Ai and Ai + 1 and equal to 300 μm and n the total number of sections analyzed in the sample.

### Immunostaining and quantification of fluorescent signal

Cryosections were defrosted at room temperature, protected from light and rehydrated in PBS. Sections were incubated 1 hour at room temperature in 5% serum, 0.25% Triton PBS before incubation with primary antibodies listed in the Table S1 overnight at 4°C. Secondary antibody incubation was performed at room temperature for 1 hour. Slides were mounted with Fluoromount-G mounting medium with DAPI (00-4959-52, Life Technologies). Slides were scanned with Zeiss Axioscan 7 and quantification of the different cell types was done using Qupath software version 0.4.0^64^.

### Flow cytometry analyses of fracture environment

#### Cell isolation

We isolated cells from the fracture environment (hematoma/callus, activated periosteum and muscle surrounded the fracture site) as described previously^8^. Tissues from uninjured tibias or from the fracture site at days 1, 3 and 5 post-injury were dissected free of fascia, tendon and fat. Tissues were minced and digested at 37°C for 2 hours in DMEM (21063029, Life Technologies) with 1% Trypsin (15090046, Life Technologies) and 1% collagenase D (11088866001, Roche). Cells in suspension were removed every 20 min and the digestion medium was replaced. After 2 hours, the cell suspension was filtered, centrifuged and resuspended for analysis.

#### Flow cytometry analysis

Digested cells were blocked with Fc-Block (anti-CD16/32, eBioscience) for 10 min on ice and stained with cell surface antibodies listed in Table S1 for 30 min, on ice and protected from light. After washing, cells were fixed and permeabilized using eBioscience Intracellular Fixation & Permeabilization Buffer Set (Invitrogen, 88_8824-00) before incubation with intracellular markers. After washes, cells were resuspended in MEMα (32561094, ThermoFischer Scientific with 10% lot-selected Fetal Bovine Serum (10270106, ThermoFischer Scientific). Analyses were performed using a BD Fortessa X20 (BD Biosciences). Data were analyzed using FlowJo v10.8.1.

### Nuclei extraction and single nuclei RNA sequencing

#### Nuclei extraction

For this study, we generated 2 datasets: a day 1 post-fracture single nuclei RNAseq dataset that we combined with datasets of the uninjured periosteum, and the periosteum and hematoma at days 3, 5 and 7 post-fracture from^4^, and a day 5 post-MTI snRNAseq dataset that we combined with the day 5 post-fracture from^4^. Nuclei extraction protocol is described in^4^. Briefly, injured tibias from 3 or 4 mice were collected and the surrounded tissues were removed. The activated periosteum was scraped and collected with the hematoma/fracture callus. Collected tissues were minced and placed 5 min in ice-cold Nuclei Buffer (NUC101, Merck) before mechanical nuclei extraction using a glass douncer. Nuclei suspension was filtered, centrifuged and resuspended in RNAse-free PBS (AM9624, ThermoFischer Scientific) with 2% Bovine Serum Albumin (A2153, Merck) and 0.2 U/µL RNAse inhibitor (3335399001, Roche). Sytox™ AADvanced™ (S10349, ThermoFischer Scientific) was added (1/200) to label nuclei and Sytox-AAD+ nuclei were sorted using Sony SH800.

#### Single nuclei RNA sequencing

The snRNA-seq libraries were generated using Chromium Single Cell Next GEM 3′ Library & Gel Bead Kit v.3.1 (10x Genomics) according to the manufacturer’s protocol. Briefly, 10 000 nuclei were loaded in the 10x Chromium Controller to generate single-nuclei gel-beads in emulsion. After reverse transcription, gel-beads in emulsion were disrupted. Barcoded complementary DNA was isolated and amplified by PCR. Following fragmentation, end repair and A-tailing, sample indexes were added during index PCR. The purified libraries were sequenced on a Novaseq (Illumina) with 28 cycles of read 1, 8 cycles of i7 index and 91 cycles of read 2. Sequencing data were processed using the Cell Ranger Count pipeline and reads were mapped on the mm10 reference mouse genome with intronic and exonic sequences.

### Single nuclei RNAseq analyses

Single-nuclei RNAseq analyses were performed using Seurat v4.1.0 ^65,66^ and Rstudio v1.4.1717.

#### Filtering and clustering

##### Periosteum and callus/hematoma

Aligned snRNAseq datasets were filtered to retain only nuclei expressing between 200 and 5000 genes and expressing less than 0,5% of mitochondrial genes and 2.5% of ribosomal genes. Contamination from myogenic cells were removed from the analyses. After filtering, the datasets of murine samples were composed of 1203 cells for uninjured datasets, 634 nuclei for day 1 post-fracture dataset, 1370 nuclei for day 3 post-fracture datasets, 1971 nuclei for day 5 post-fracture dataset and 997 nuclei for day 7 post-fracture dataset. The datasets were merged and scaled on mitochondrial and ribosomal content and cell cycle. Clustering was performed using the first 20 principal components and a resolution of 1.7. Immune cells clusters from the integration were isolated to perform subset analysis and was reclustered using the first 10 principal components and a resolution of 1,7. SSPC, fibrogenic, chondrogenic and osteogenic clusters from the integration were isolated to perform subset analysis. The subset was reclustered using the first 20 principal components and a resolution of 0.5.

For the day 5 post-MTI dataset, we filtered nuclei expressing between 200 and 5000 genes and expressing less than 5% of mitochondrial genes and 2% of ribosomal genes. After filtering and removal of myogenic cells, we obtained X nuclei. We integrated the dataset with the day 5 post-fracture dataset, scaled on mitochondrial and ribosomal content and clustered using the first 30 principal components and a resolution of 1. For subset analyses, we used the 20 first component and a resolution of 1.5 for the immune subset and the 15 first component and a resolution of 0.5 for the IIFC subset.

##### Skeletal muscle

Datasets from ^8^ were used for these analyses (GSE195940). These datasets correspond to sorted GFP+ cells isolated from the skeletal muscle surrounding the tibia of *Prx1^Cre^; R26^mTmG^* mice without injury and at days 3 and 5 post-fracture and MTI. Cells expressing between 1000 and 8000 genes and <20% mitochondrial genes were retained for analysis. Pericytes and tenocytes were removed from the analyses. Integrated analysis of days 0, 3 and 5, post-fracture was regressed using mitochondrial content, the number of genes detected within each cell, and performed using top 2000 features and the 30 first components with a resolution set at 1.3. Integrated dataset of days 0, 3 and 5 post-fracture and MTI was regressed using mitochondrial and ribosomal content and then performed using top 2000 features and the 25 first components with a resolution set at 1.1.

#### Pseudotime analysis

Monocle3 v1.0.0 was used for pseudotime analysis ^67^. Single-cell trajectories were determined using monocle3 default parameters. The starting point of the pseudotime trajectory was determined as the cells from the uninjured dataset with the highest expression of stem/progenitor marker genes *(Ly6a, Cd34, Dpp4, Pi16)*.

#### Lineage score

Lineage score was calculated by the mean of the expression of specific markers from the literature listed in Table S2.

#### Single cell regulatory network inference using SCENIC

Single cell regulatory network inference and clustering (SCENIC) ^38^ was used to infer transcription factor (TF) networks active in the subsets of IIFCs. Analysis was performed using recommended parameters using the packages SCENIC v1.3.1, AUCell v1.16.0, and RcisTarget v1.14 and the motif databases RcisTarget and GRNboost.

#### Cell-cell interaction using CellChat and Connectome

Cell communication analysis was performed using the R package CellChat ^37^ and Connectome ^68^ with default parameters on the complete fracture combined dataset and on the subset of immune cells.

### Statistical analysis

Data are presented as mean ± s.d. and were obtained from at least two independent experiments. Statistical significance was determined with a two-sided Mann–Whitney test and reported from GraphPad Prism v9.0.2. Differences were considered to be significant when p< 0.05, and reported as * when p<0.05 and ** when p<0.01.

## Acknowledgements

We thank A. Guigan, O. Ruckebusch and A. Henry from the Flow Cytometry platforms of IMRB, L. Slimani and K. Henri from Life Imaging Facility of Paris Cité University (Plateforme Imagerie du Vivant “Micro-CT platform”), the staff from the IMRB genomic and bioinformatic platforms for advice and technical assistance. We thank C. Goachet, V. Bretegnier, G.Chemin, E. Rohart and S. Lakhlifi for technical assistance or advice.

## Author contribution

<u>Conceptualization:</u> Y.H., S.P., A.J., C.C. <u>Methodology:</u> Y.H., S.P., C.C. <u>Formal Analysis:</u> Y.H., S.P. <u>Investigation:</u> Y.H., S.P., A.J., M.E., J.V., B.G., <u>Resources:</u> C.G. <u>Writing – Original Draft:</u> Y.H., S.P., C.C. <u>Writing – Review & Editing:</u> A.J., M.E. <u>Visualization:</u> Y.H., S.P. <u>Supervision:</u> C.C. <u>Funding Acquisition:</u> C.C.

## Data and materials availability

The single-nuclei RNAseq dataset generated for this study will be deposited on GEO and will be made publicly available as of the date of publication. Single-nuclei RNAseq datasets from Perrin *et al.* are deposit on GEO (GSE234451). This paper does not report original code.

