## Supplemental file for "Multimodal analyses of immune cells during bone repair identify macrophages as a therapeutic target in musculoskeletal trauma"

Supplementary material

**Figure S1**

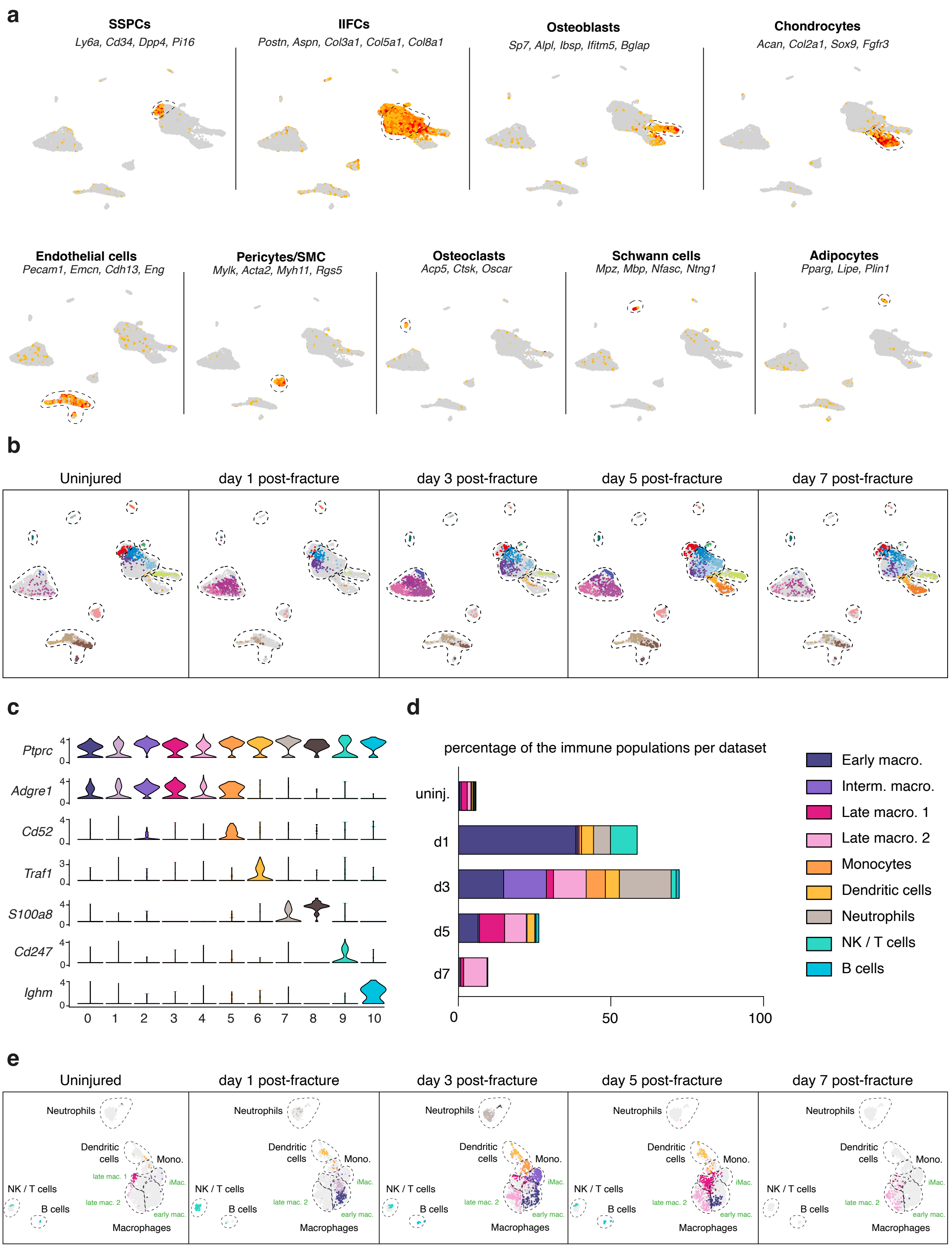


### Supplemental Figure 1: Immune cell atlas of bone regeneration

**a.** Feature plots of the lineage score of the different cell populations in the combined fracture datasets. **b.** UMAP projection of the cells from the different time points in the integrated dataset. **c.** Violin plots of the expression of marker genes of the different immune cell populations. **d.** Percentage of cells in the different immune cell populations per time point. **e.** UMAP projection of the cells from the different time points in the integrated dataset.

Figure S2

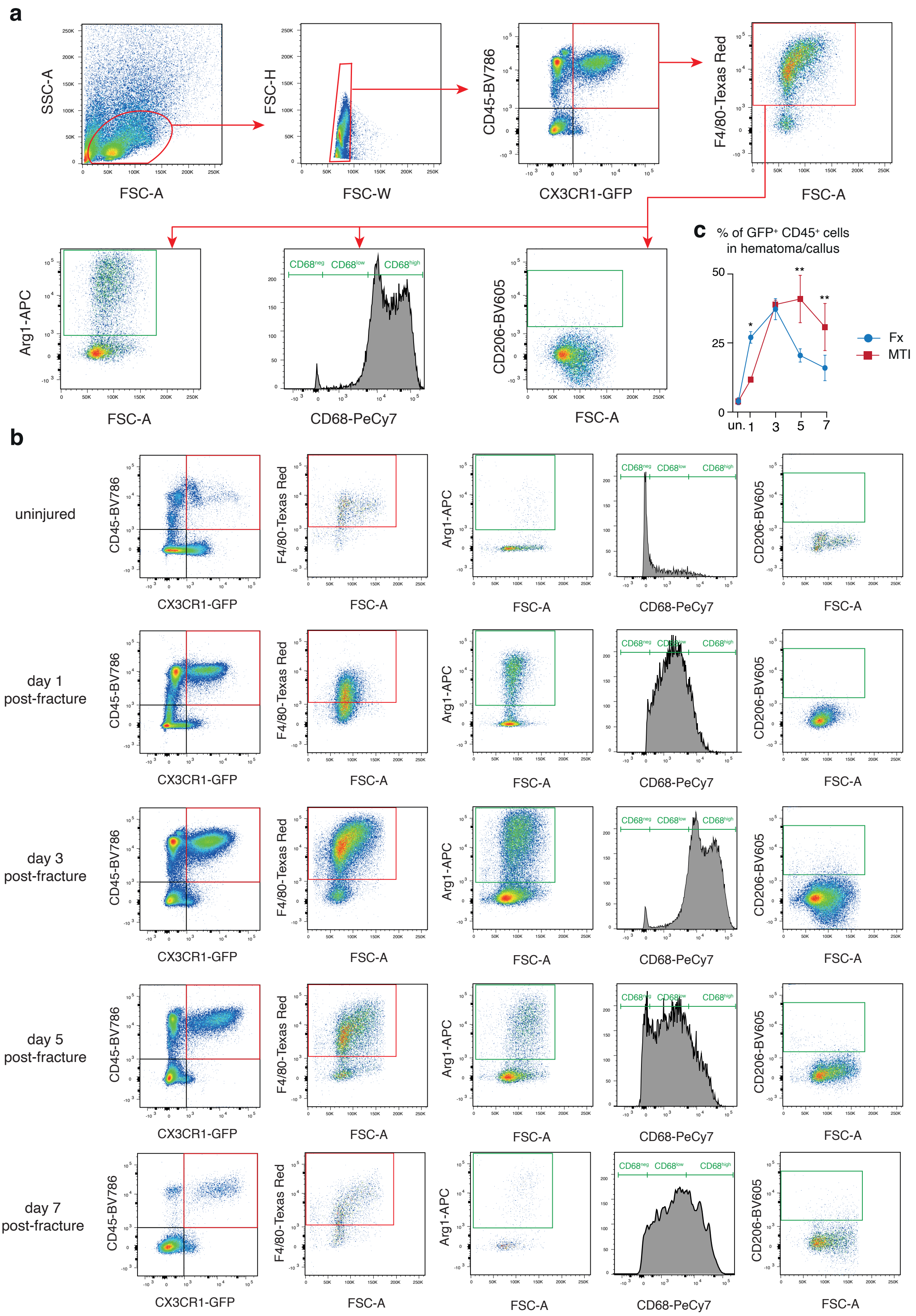


**Supplemental Figure 2: Flow cytometry analyses of the macrophage dynamics after bone fracture**

**a.** Gating strategy of the flow cytometry analyses of macrophages in the fracture environment. **b**. Flow cytometry analyses of macrophages in uninjured periosteum and muscle and in the fracture environment at days 1, 3, 5 and 7 post-fracture. **c**. Percentage of CD45+ GFP+ cells in uninjured periosteum/muscle and in the fracture environment at 1-, 3-, 5- and 7-days post-fracture of *Cx3cr1*$^{GFP}$ mice.

**Figure S3**

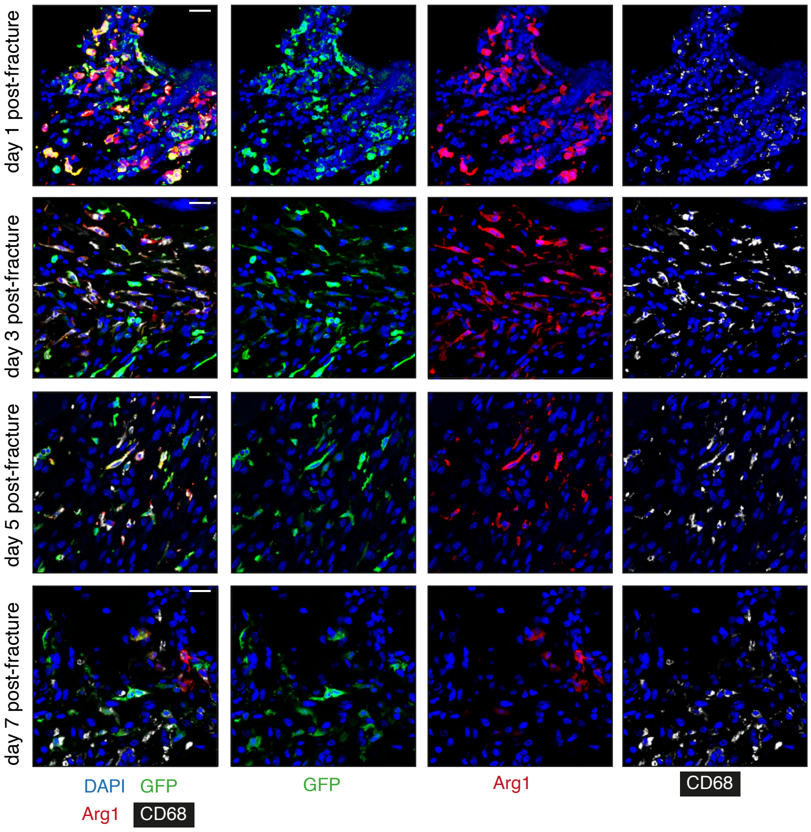


**Supplemental Figure 3: Macrophages express both CD68 and Arg1 during the early steps of bone repair**

Co-immunostaining of Arg1 and CD68 in hematoma and callus fibrosis at days 1, 3, 5 and 7 post-fracture of *Cx3cr1*$^{GFP}$ *mice*. Scale bars: 25 µm

**Figure S4**

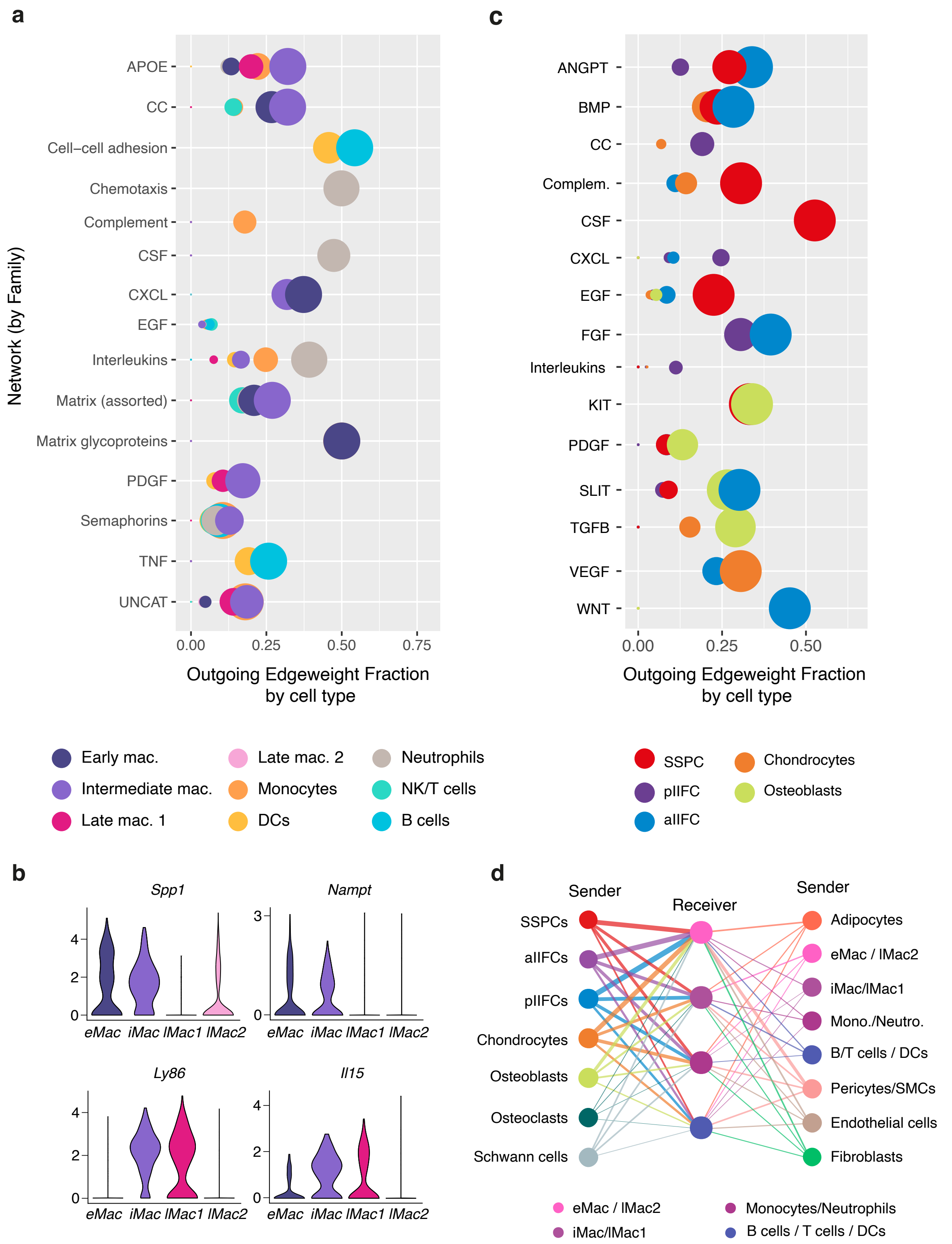


**Supplemental Figure 4: Paracrine interactions between immune cells and SSPCs during bone repair**

**a.** Centrality analyses of the outgoing signals from the different immune cell populations after injury. **b.** Feature plots showing the expression of *Spp1* and *Nampt* by early macrophages (eMac) and intermediate macrophages (iMac) and the expression of *Ly86* and *Il15* by both iMac and late macrophages 1 (lMac1). **c.** Centrality analyses of the outgoing signals from the SSPCs, IIFCs, chondrocytes and osteoblasts after fracture. **d.** Interaction plot showing the interaction strengths from the different populations of the single nuclei dataset to the immune cells. The level of estimated interaction is displayed by the line thickness.

**Figure S5**

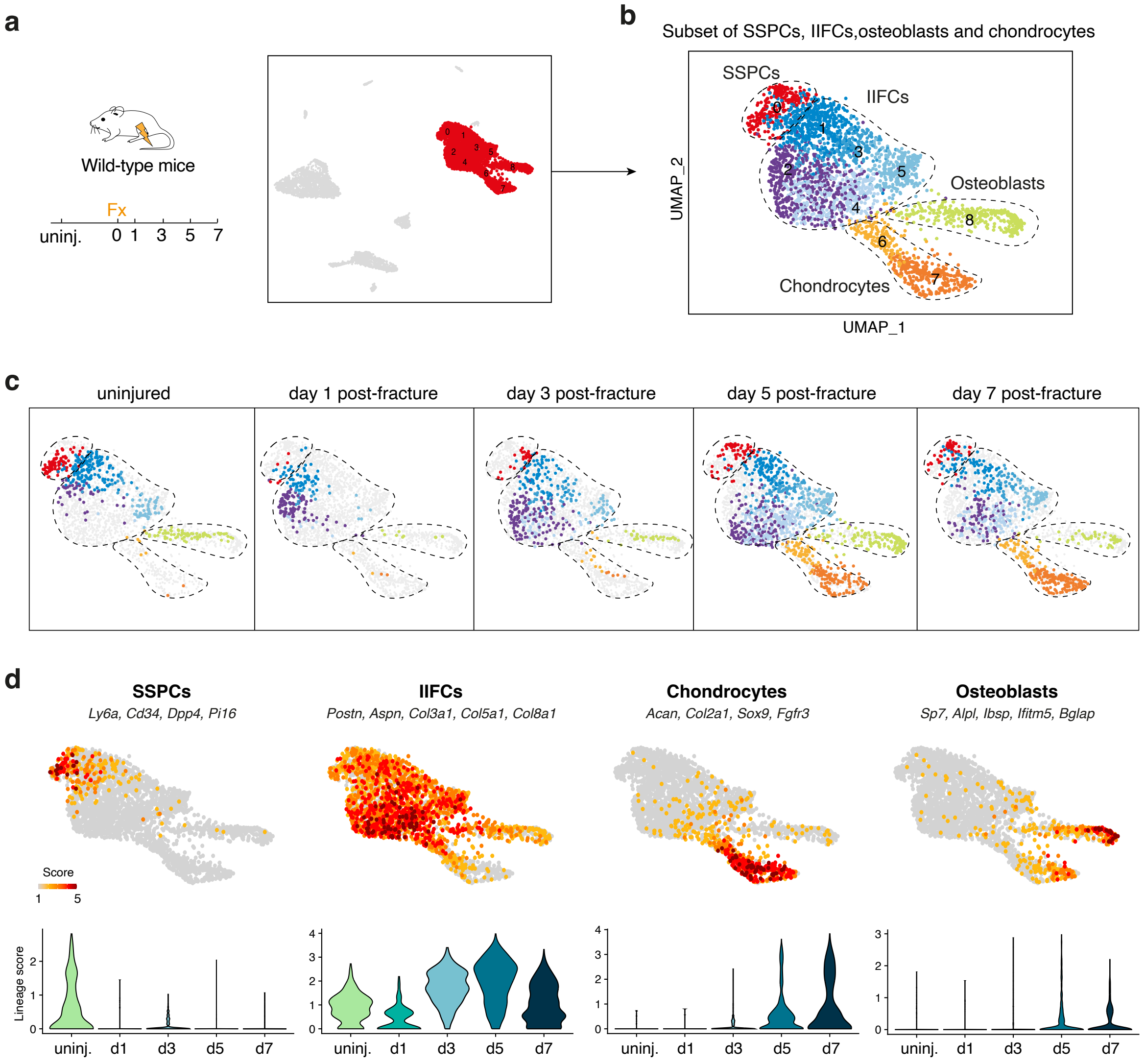


**Supplemental Figure 5: Periosteal SSPCs differentiate via a common fibrogenic stage prior to osteogenesis or chondrogenesis**

**a.** Experimental design. SSPCs, injury-induced fibrogenic cells (IIFCs), chondrocytes and osteoblasts from integrated uninjured, day 1, day 3, day 5 and day 7 post-fracture samples were extracted for a subset analysis. **b**. UMAP projection of color-coded clustering of the subset dataset. The four populations are delimited by black dashed lines. **c.** UMAP projection of the cells from the different time points in the subset of SSPCs, IIFCs, chondrocytes and osteoblasts. **d.** (top) Feature plots of the stem/progenitor, IIFC, chondrogenic and osteogenic lineage scores. (bottom) Violin plots of the lineage score per time point.

**Figure S6**

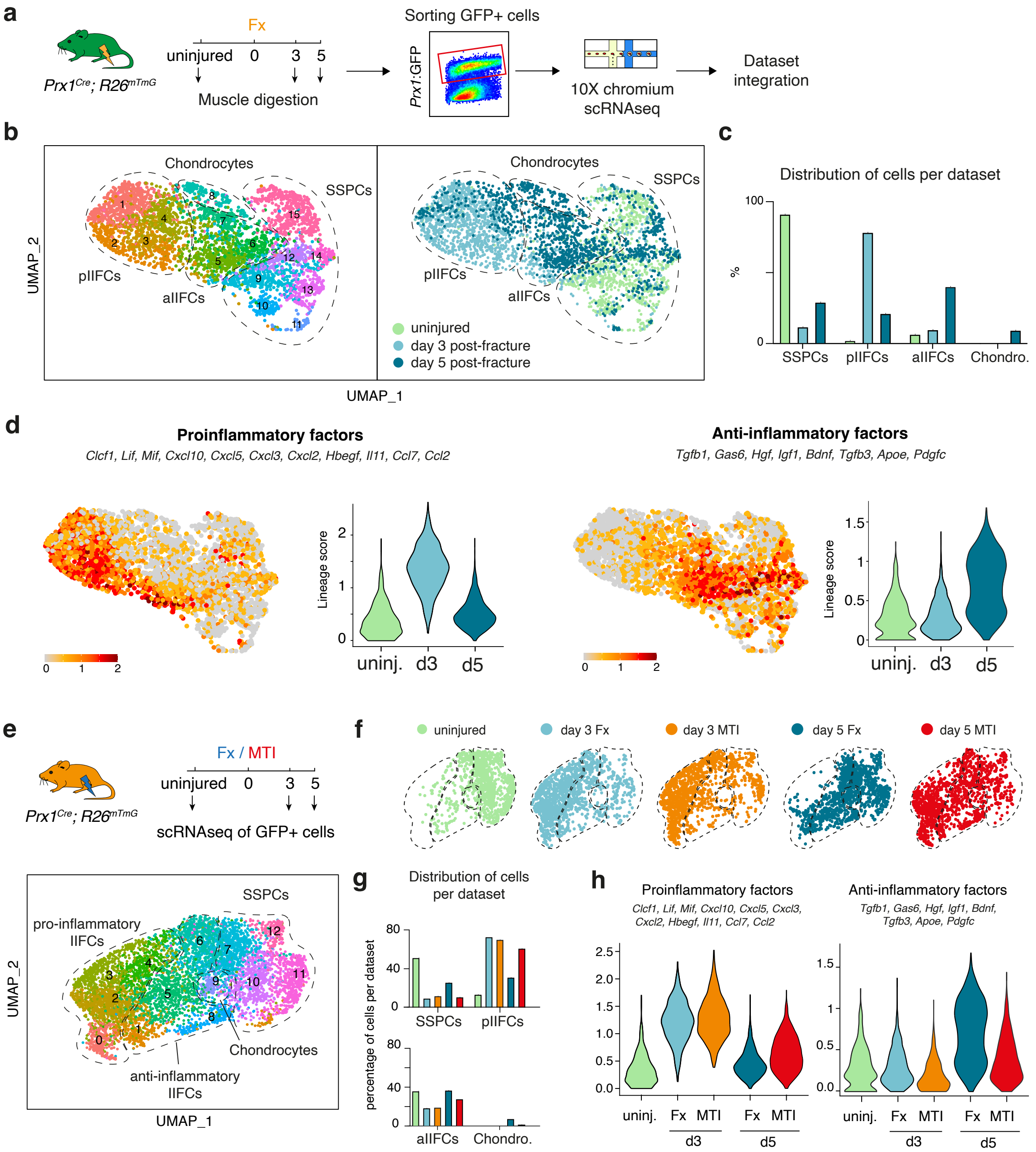


**Supplemental Figure 6: The immune response of skeletal muscle SSPCs is altered after musculoskeletal trauma**

**a.** Experimental design. Prx1-derived skeletal muscle cells were isolated from hindlimb skeletal muscles and sorted based on GFP-expression prior to scRNAseq[1]. **b.** Left, UMAP projection of color-coded clustering of Prx1-derived cells reveals 15 clusters defining 4 distinct populations (limited by a black dashed line). Right, UMAP projection of the integrated datasets generated from uninjured muscle and from muscles at day 3 and day 5 post-fracture, colored by time point. **c.** Percentage of cells per population at each time point. **d.** Feature plots and violin plots of the expression of pro-inflammatory and anti-inflammatory factors per time point. **e.** (top) Experimental design of scRNAseq experiment. Prx1-derived skeletal muscle cells were isolated from uninjured muscle and from muscles at day 3 and day 5 post-fracture or MTI and sorted based on GFP-expression prior to scRNAseq [1]. (bottom) UMAP projection of color-coded clustering of Prx1-derived SSPCs and derivatives reveals 13 clusters defining 4 distinct populations (limited by a black dashed line). **f.** UMAP projection of the integrated scRNAseq dataset separated by dataset. **g.** Distribution of cells per population per time point. **h.** Violin plots of the expression of pro-inflammatory and anti-inflammatory factors per dataset.

**Figure S7**

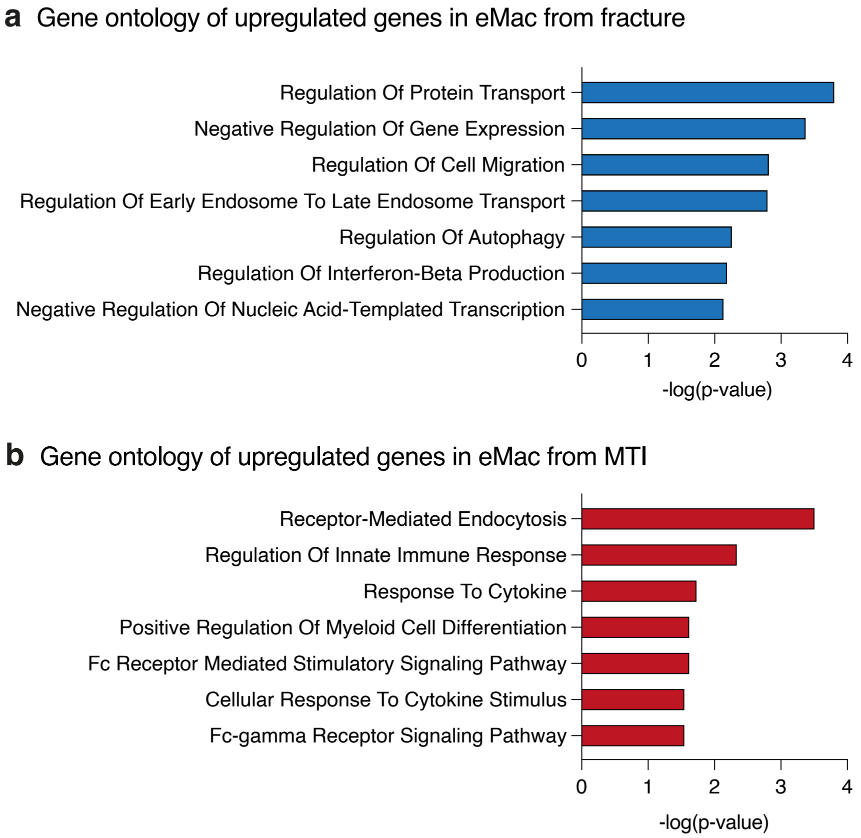


**Supplemental Figure 7: Prolonged immune activity in early macrophages in MTI**

Gene ontology analyses of upregulated genes in eMac at day 5 post-injury from the fracture dataset (**a**) and MTI dataset (**b**).

**Figure S8**

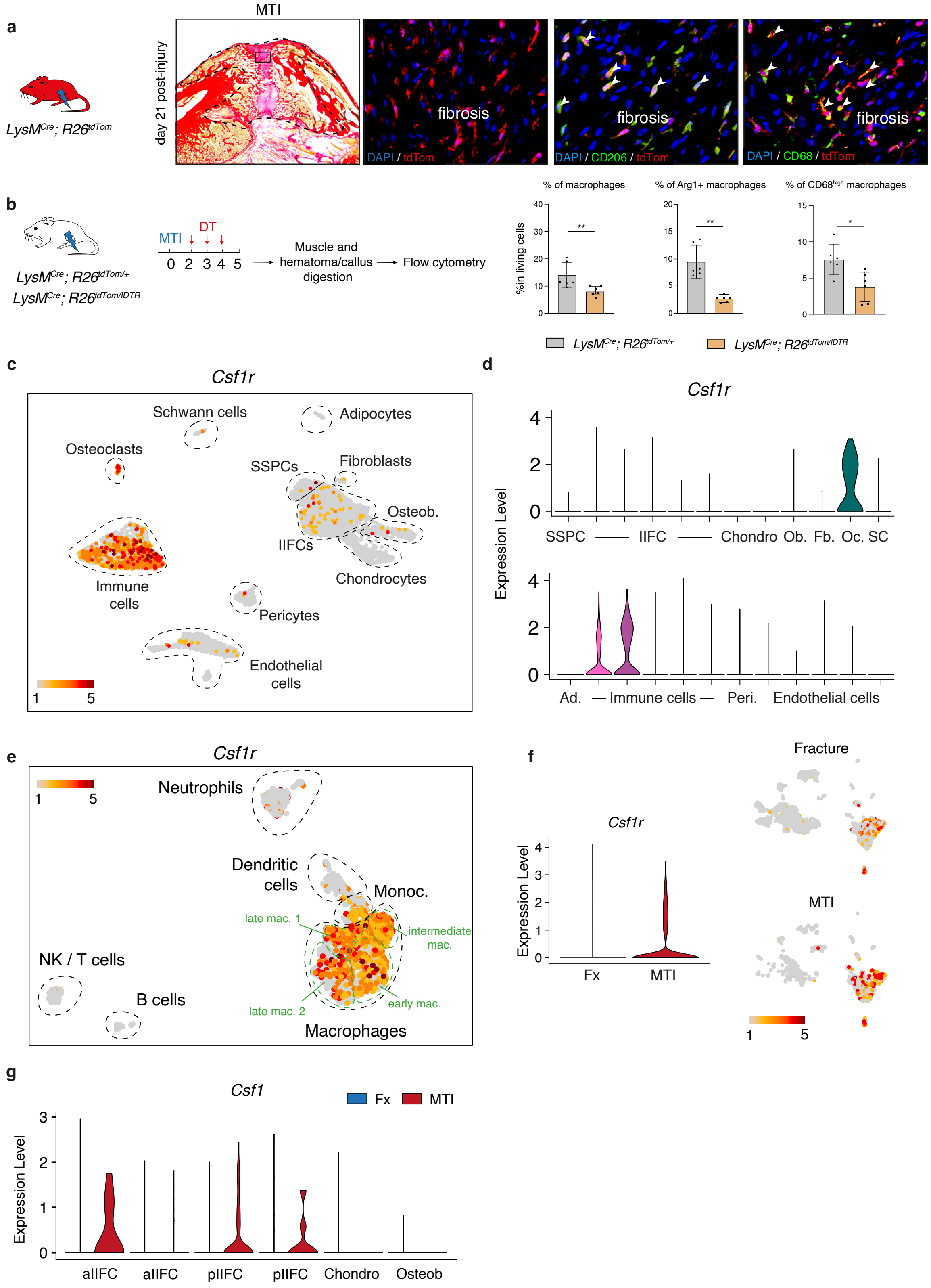


**Supplemental Figure 8: Macrophages in MTI-induced fibrosis can be targeted in the *LysM$^{Cre}$* mouse model and express *Csf1r***

**a.** Callus section of 21 days post-MTI in *LysM*$^{Cre}$*; R26*$^{tdTom}$ mice shows the presence of tdTom+ cells in the fibrotic tissue. Co-immunofluorescence of CD68 and CD206 shows that tdTom+ cells are CD68 and CD206 positive macrophages. **b.** Experimental design of macrophage depletion in *LysM*$^{Cre}$*; R26*$^{tdTom:/iDTR}$ mice and assessment of depletion efficiency by flow cytometry. Mice received DT injections at days 2, 3 and 4 post-MTI and cells were isolated from the fracture environment (hematoma/callus and muscle) at day 5. *LysM*$^{Cre}$*; R26*$^{tdTom/iDTR}$ mice showed a reduction in the percentage of macrophages (CD45+ Tomato+ F4/80+ cells) and the percentage of Arg1+ and CD68$^{High}$ macrophages (n=6 per group). **c.** Feature plot of *Csf1r* expression in the complete integrated snRNAseq dataset from the periosteum and hematoma/callus at days 1, 3, 5, and 7 post-fracture. **d.** Violin plot of *Csf1r* expression in the different clusters from the integrated fracture dataset. **e.** Feature plot of *Csf1r* expression in the subset of immune cells. **f.** Violin plot and feature plots of integrated dataset separated by injury type of *Csf1r* expression in the day 5 post-fracture and post-MTI snRNAseq datasets. **g.** Feature plot of *Csf1* expression by IIFCs, chondrocytes and osteoblasts at day 5 post-fracture (blue) and MTI (red).

| Antibody type | Use | Antigen | Reference |
|---|---|---|---|
| Primary | IF | rat monoclonal to mouse CD68 | Biolegend, 137002 |
| Primary | IF | rabbit polyclonal to mouse Arginase 1 | Abcam, ab91279 |
| Primary | IF | rabbit polyclonal to mouse CD206 | Abcam, ab64693 |
| Primary | IF | chicken polyclonal to GFP | Abcam, ab13970 |
| Secondary | IF | Alexa Fluor 647 Goat anti-Rat IgG | Invitrogen, A21247 |
| Secondary | IF | Alexa Fluor 555 Goat anti-Rabbit IgG | Invitrogen, A21428 |
| Secondary | IF | Alexa Fluor 488 Goat-anti Chicken IgG | Invitrogen, A11039 |
| Primary | FC | anti-CD45 BV786 | BD Bioscience, 564225 |
| Primary | FC | anti-F4/80 | BD Bioscience, 565613 |
| Primary | FC | anti-Arg1 APC | Invitrogen, 17-3697-82 |
| Primary | FC | anti-CD206 BV605 | Biolegend, 141723 |
| Primary | FC | anti-CD68 Pe-Cy7 | Invitrogen, 25-0681-82 |
| Primary | FC | anti-F4/80 | BD Bioscience, 567201 |

**Supplementary Table 1: List of antibodies used for this study.**

IF: immunofluorescence. FC: flow cytometry.

| Score | Genes |
|---|---|
| SSPCs | *Ly6a, Cd34, Dpp4, Pi16* |
| IIFCs | *Postn, Aspn, Col3a1, Col5a1, Col8a1* |
| Osteoblasts | *Sp7, Alpl, Ibsp, Ifitm5, Bglap* |
| Chondrocytes | *Acan, Col2a1, Sox9, Fgfr3* |
| Immune cells | *Ptprc, Adgre1, Cd8, Cx3cr1* |
| Endothelial cells | *Pecam1, Emcn, Cdh13, Eng* |
| Pericytes/SMC | *Mylk, Acta2, Myh11, Rgs5* |
| Osteoclasts | *Acp5, Ctsk, Oscar* |
| Schwann cells | *Mpz, Mbp, Nfasc, Ntng1* |
| Adipocytes | *Pparg, Lipe, Plin1* |
| Macrophage pro-inflammatory secretome | *Cxcl2, Ccl7, Ccl2, Ccl9, Thbs1* |
| Macrophage pro-repair secretome | *Apoe, Hgf, Cxcl16, Tgfb1, Pf1, Pltp, Sema4d, Pros1* |
| Macrophage anti-inflammatory secretome | *Igf1, Gas6, Pdgfc* |
| Periosteal IIFC pro-inflammatory factors | *Cxcl2, Cxcl5, Ccl2, Ccl5, Ccl7, Clcf1, Csf1, Lif, Mif, Cit* |
| Periosteal IIFC anti-inflammatory factors | *Igf1, Bdnf, Tgfb3, Gas6, Hgf, Inhba* |
| Muscle IIFC pro-inflammatory factors | *Clc1, Lif, Mif, Cxcl10, Cxcl5, Cxcl3, Cxcl2, Hbegf, Il11, Ccl7, Ccl2* |
| Muscle IIFC anti-inflammatory factors | *Tgfb1, Gas6, Hgf, Igf1, Bdnf, Tgfb3, Apoe, Pdgfc* |

**Supplementary table 2: Lists of genes used for lineage/secretome score analyses.**
